# Head Movements Control the Activity of Primary Visual Cortex in a Luminance Dependent Manner

**DOI:** 10.1101/2020.01.20.913160

**Authors:** Guy Bouvier, Yuta Senzai, Massimo Scanziani

## Abstract

The vestibular system broadcasts head-movement related signals to sensory areas throughout the brain, including visual cortex. These signals are crucial for the brain’s ability to assess whether motion of the visual scene results from the animal’s head-movements. How head-movements impact visual cortical circuits remains, however, poorly understood. Here, we discover that ambient luminance profoundly transforms how mouse primary visual cortex (V1) processes head-movements. While in darkness, head movements result in an overall suppression of neuronal activity, in ambient light the same head movements trigger excitation across all cortical layers. This light-dependent switch in how V1 processes head-movements is controlled by somatostatin expressing (SOM) inhibitory neurons, which are excited by head movements in dark but not in light. This study thus reveals a light-dependent switch in the response of V1 to head-movements and identifies a circuit in which SOM cells are key integrators of vestibular and luminance signals.

## Introduction

Primary sensory areas of the mammalian cortex are each dedicated to one sensory modality defined by the sensory organs they receive input from. The vestibular organs detect angular rotation and linear acceleration of the head, and transmit these signals to the brain^1, 2^. However, unlike most other senses, no primary cortical area is dedicated to the processing of vestibular signals. Instead, head-movement related signals are broadcast across cortical areas dedicated to distinct sensory modalities^3–7^. The integration of vestibular signals with other sensory modalities likely allows the brain to assess whether a given sensory stimulus results from the animal’s head-movement in the environment rather than from a change in the sensory environment itself. Primary visual cortex (V1) of the mouse is one of those sensory areas which, in addition to receiving input from their main sensory organ, the retina, also receives head-movement related signals from vestibular organs^4^. While we have extensive knowledge of how cortical circuits in V1 process visual stimuli, we still have only a rudimentary understanding of how those same primary visual circuits process vestibular signals.

Prior work on how vestibular stimuli impact activity in visual cortex has led to different observations. Studies in humans indicate that vestibular stimulation reduces basal activity in visual cortex, implying an overall suppressive effect of the vestibular system on this structure^8, 9^. However, more recent work in mice has shown that head movements increase activity in layer 6 pyramidal neurons of V1, instead implying an excitatory effect, at least in this layer^4^. Other studies in primates and carnivores have shown more heterogeneous and complex effects of vestibular stimulation on visual cortex activity, depending on the type of vestibular stimuli and the properties of the visual stimuli presented concomitantly to the vestibular stimulus^5, 6, 10, 11^. The impact of vestibular stimuli on V1 may also depend on ambient luminance. In fact, V1 has been shown to adapt to changes in luminance through sustained changes in its basal activity. Thus, depending on ambient luminance, and hence on its basal activity, V1 may differentially process incoming vestibular information. In other words, the same head movements occurring under different ambient luminance levels, as an animal explores its environment, may differently impact V1.

Here, using extracellular recordings in head-fixed and freely moving mice, we show that head movements control V1 activity in a luminance dependent manner. In the dark, head movements exert an overall suppressive action on neuronal activity in V1 while in light, the same head movements robustly shifted cortical activity toward excitation across all layers. This light-mediated shift in vestibular responses impacts both pyramidal cells and parvalbumin-expressing (PV) inhibitory cells in a similar manner, while exerting an opposite effect on somatostatin expressing (SOM) inhibitory cells. Finally, we show that genetic ablation of SOM cells strongly reduces both the suppression of V1 in response to head movements in the dark and the shift toward excitation in light. This study reveals a light-dependent vestibular impact on V1 and identifies a circuit in which SOM cells are key integrators of vestibular and luminance signals.

## Results

### Suppression of V1 by head movements in the dark

How does V1 respond to head movements? To control the velocity and amplitude of head movements, we fixed the head of awake mice in the center of a servo-controlled table enabling the rotation of the animal along the horizontal plane (50° rotation; 80°/s peak velocity; unless stated otherwise Fig. 1a). Using extracellular linear probes, we recorded from neurons in V1 in the left hemisphere across all cortical layers and prevented visual stimuli to contaminate the recorded responses by performing the experiments in the dark.

**Figure 1.**
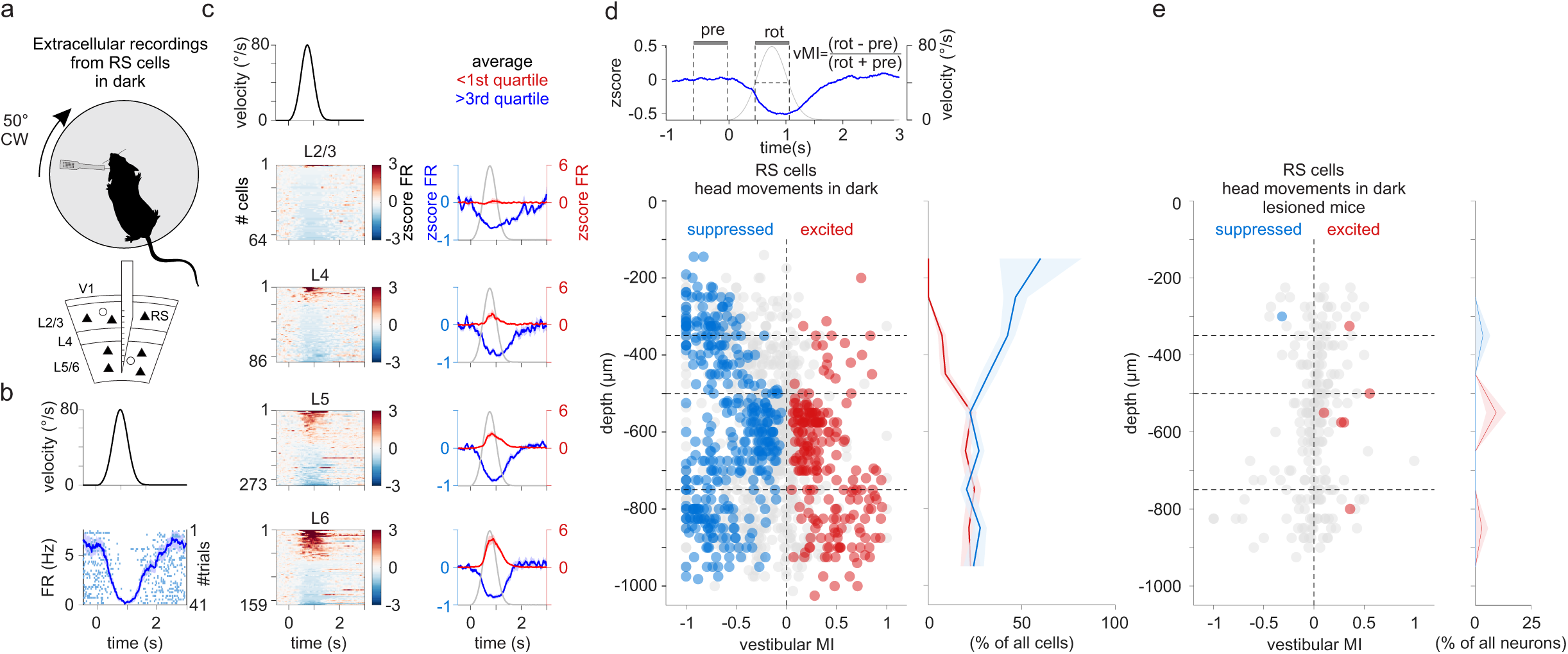
Suppression of V1 by head movements in dark. **(a)** Experimental configuration: Top: An extracellular linear probe in the left primary visual cortex (V1) of a head-fixed, awake mouse records the response to clockwise (CW) rotations of the table in dark. Bottom: The linear probe spanned all cortical layers. RS: regular spiking cell. **(b)** Top: Velocity profile of the rotating table (CW rotation). Bottom: Example RS cell located 550 μm below the pial surface. Raster plot and averaged peristimulus time histogram (PSTH) are superimposed. Shading is SEM. **(c)** Summary of 582 RS cells significantly modulated cells subdivided according to cortical depth (from the pia). Left panels: Velocity profile of the rotating table (CW rotation; top panel) and heat maps of the z-score of the firing rate (zscore FR) of individual RS cells during rotations. Within each panel, cells are sorted by the peak z-score. Right panels: Blue and red traces are the average z-scores of all cells whose z-score was smaller or larger than the first or third quartile, respectively. Shaded area is SEM. The gray trace is the velocity profile (CW rotation). **(d)** Top left: The time windows used to compute the vestibular modulation index (vMI; blue trace: average of all significantly suppressed RS cells; gray trace: velocity profile). Bottom left: vMI for CW rotations of individual RS cells plotted against cortical depth. Blue, red and gray circles are suppressed, excited and non-significantly modulated RS cells, respectively (superficial layers vMI: −0.45±0.02; n = 353 cells; Deep layers vMI: −0.15±0.01; n = 978 cells; 36 recordings from 30 mice). Horizontal dotted lines indicate approximate layer borders. Right: Percentage of suppressed (blue) and excited (red) RS cells plotted against to cortical depth (superficial layers percentages: 32±4% suppressed; 6±1% excited, Deep layers percentages: 25±2% suppressed; 22±2% excited). Shading is SEM. **(e)** As d but for vestibular lesioned mice (2.7% of RS cells modulated by rotation, 7 out of 253 units, n = 3 mice).

The firing rate of most V1 neurons (62%; 928 out of 1502 cells; n = 30 animals) was modulated by either clockwise (CW; contraversive) or counterclockwise (CCW; ipsiversive) rotations of the table (Fig. 1b, Supplementary Fig. 1a-c). The time-course of this modulation approximated the velocity profile of the rotating table (Supplementary Fig. 1c) and 40% of the modulated neurons showed a significant difference in the response to CW or CCW rotations (Supplementary Fig. 1b). We quantified the response of V1 neurons by computing the vestibular modulation index (vMI, Fig. 1d, top panel). Positive or negative vMIs indicate, respectively, a head movement mediated increase (excitation) or decrease (suppression) in activity relative to baseline. Strikingly, both CW and CCW rotations of the table led to a strong overall suppression of neuronal activity in V1, particularly in the superficial layers (layer 2/3 and layer 4; Fig. 1c,d and Supplementary Fig. 1a). In contrast, neurons in deep layers were approximately equally distributed between those that were suppressed and those that were excited by the rotation (Fig. 1c,d and Supplementary Fig. 1a). This was the case for both regular-spiking (RS; putative excitatory neurons, Fig. 1) and fast-spiking cells (FS; putative parvalbumin-expressing inhibitory cells, Supplementary Fig. 2). Furthermore, the overall suppressive impact of head movements on V1 activity was observed independent of the specific velocity profile or peak velocity used to rotate the table (Supplementary Fig. 1d).

The response of V1 neurons to the rotation of the table was due to the activation of vestibular organs because bilateral vestibular lesions abolished the response (Fig. 1e). Moreover, even though V1 can respond to sound^12, 13^, the response to table rotations was not due to the sound of the servo motor because detaching the table from the motor, hence preserving the sound without triggering rotation, did not elicit any response (3.8%, 13 out of 341 cells, n = 3 mice, data not shown).

Taken together, these data show that head movements control the activity of large fractions of V1 neurons across all layers and exert an overall suppressive impact, especially in superficial layers.

### Excitation of V1 by head movements in light

Because V1 basal activity is regulated by ambient luminance^14–17^, we compared the response of V1 to head movements in dark and light conditions. To avoid contaminating V1 responses to head movements in light with responses to the visual environment, we stitched both eyelids and placed a light diffuser in front of the light source to achieve a homogeneous illumination of the eye (Fig. 2a, see methods). We alternated rotations in light and dark conditions using the same velocity profile described above.

**Figure 2.**
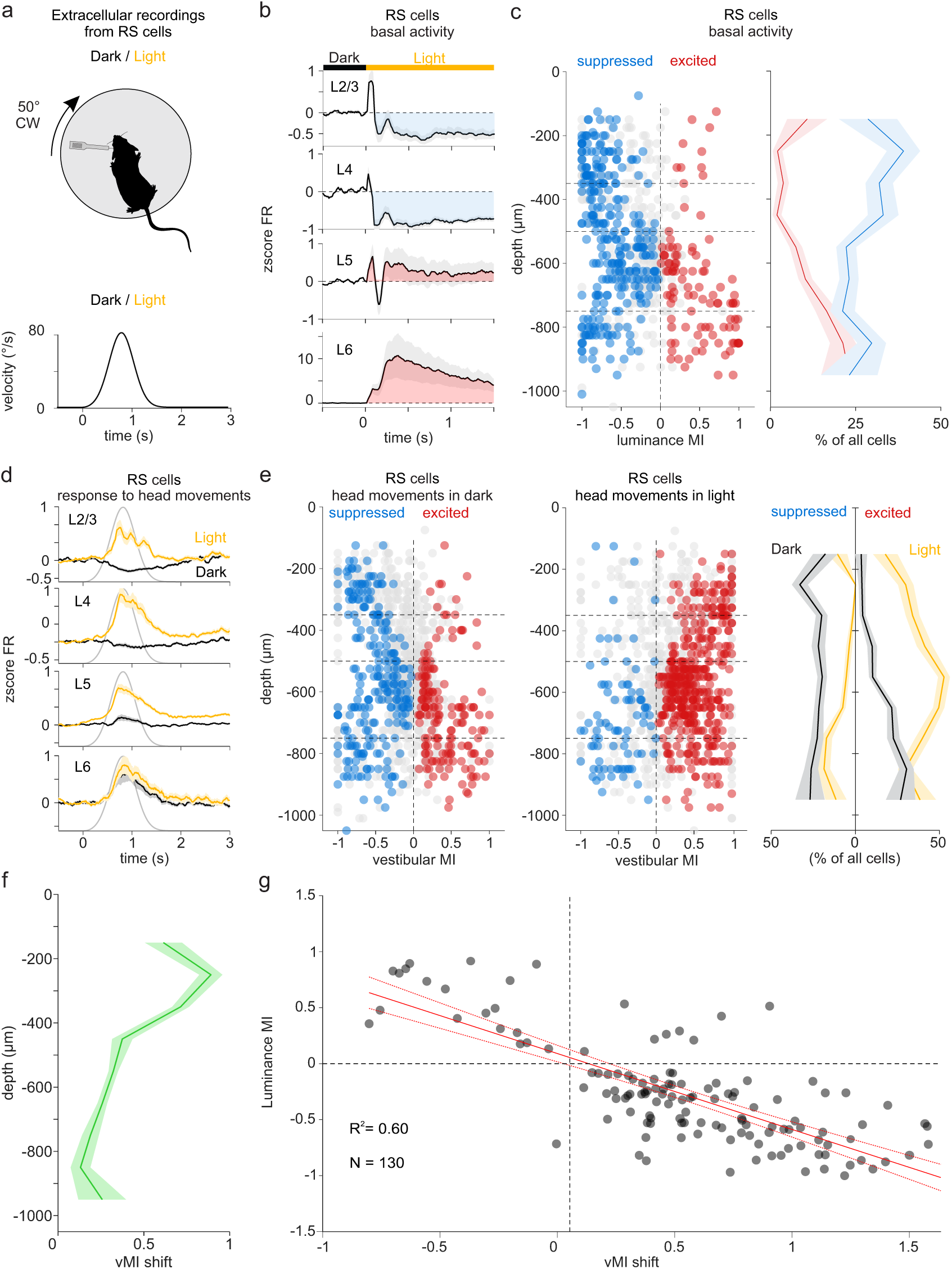
Excitation of V1 by head movements in light. **(a)** Experimental configuration: Top: As in Fig. 1a but this time rotations are alternated between dark and light. Bottom: Velocity profile of the rotation. **(b)** Averaged z-score of the firing rate (zscore FR) of RS cells in response to light onset (measured between 0.5 s and 1s). Each average is contributed by all the cells recorded within the layer indicated in the panel, independently of whether they were excited, suppressed on non-significantly modulated by light onset. Gray shading is SEM. Blue or red shaded area indicate average suppression or facilitation by light, respectively. Horizontal bar indicates time of dark/light transition (time 0). **(c)** Luminance modulation index (lumMI) of individual RS cells plotted against cortical depth. Blue, red and gray circles represent RS cells whose basal activity is suppressed, excited and non-significantly modulated by light onset, respectively. Superficial layers lumMI: −0.58±0.02; Deep layers lumMI: 0.17±0.018). Horizontal dotted lines indicate approximate layer borders. Right: percentage of suppressed (blue) and excited (red) RS cells plotted against cortical depth in response to light onset. Shading is SEM. **(d)** Averaged z-score of the firing rate (zscore FR) of RS cells during CW rotations in dark (black traces) and in light (yellow traces). Each average is contributed by all the cells recorded within the layer indicated in the panel, independently of whether they were excited, suppressed on non-significantly modulated by the rotation. Shading is SEM. Gray trace is velocity profile. **(e)** Vestibular modulation index (vMI) for CW rotations of individual RS cells plotted against cortical depth. Blue, red and gray circles are suppressed, excited and non-significantly modulated RS cells, respectively. Same cells as in C. Left: vMI of RS cells in dark; Middle, vMI of the same RS cells in light. For RS cells recorded in superficial layers, the vMI shifted from −0.30±0.02 in dark to 0.36±0.03 in light (P = 6.1e-43; n = 373 cells). In deep layers, the vMI of RS cells shifted from −0.08±0.016 in dark to 0.17±0.02 in light (P = 1.5e-33; n = 701 cells18 recordings from 12 mice). Horizontal dotted lines indicate approximate layer borders. Right: percentage of suppressed and excited RS cells plotted against cortical depth in dark (black) and light (yellow). Shading is SEM. Note the shift towards positive vMI in light. **(f)** vMI shift (vMI in light - vMI in dark) plotted against to the cortical depth. Shading is SEM. Positive vMI shift indicate facilitation of the response to head movements in light as compared to dark. **(g)** vMI shift plotted against lumMI. Included are only RS cells with a significant lumMI and a significant vMI shift (n = 130 cells). Continuous red line: linear regression (R^2^ = 0.60). Dotted red lines: confidence interval of linear regression. Note inverse relationship between vMI shift and lumMI.

Consistent with previous reports^16, 17^, light itself, i.e. prior to table rotation, had an overall suppressive impact on the basal activity of V1 neurons, especially in superficial layers (Fig. 2b,c and Supplementary Fig. 3f-h), as captured by the luminance modulation index (lumMI; positive or negative lumMIs indicate a light mediated increase or a decrease in basal activity relative to dark, respectively).

In striking contrast to the suppressive impact of head movements in the dark, head movements in the light resulted in an overall excitation of V1 cells (Fig. 2d-f and Supplementary Fig. 3d-f). Not only did light increase the average vMI across all layers, but cells in superficial layers shifted from a net average suppression in the dark to a net average excitation by head movements in light (Fig. 2d-f and Supplementary Fig. 4a-c). Despite this strong shift towards excitation, the preference of individual neurons for CW or CCW rotations remained essentially unaltered (Supplementary Fig. 4d). Consistent with the population average data, the response to head movements of most individual cells shifted towards positive vMIs in light as compared to dark (Supplementary Fig. 4a-c). Interestingly, this shift towards positive vMI depended on the impact of light on the cell’s basal activity: the larger the light mediated suppression of basal activity, the larger the positive shift of the vMI. This was the case for both RS and FS cells (Fig. 2g; Supplementary Fig. 3g). Hereinafter, we refer to the positive shift of vMI from dark to light as facilitation.

These data thus show that head movements control the activity of V1 neurons in a luminance dependent manner. While head movements mainly suppress V1 neurons in the dark, the same head movements in light excite V1 neurons.

### Excitation of layer 5 SOM cells by head movements in dark but not in light

What is the source of V1 suppression in response to head movements in the dark and how is this suppression reduced or even completely removed in light? An inhibitory neuron who is excited by head movements in the dark, thereby suppressing its targets, but is no longer excited by the same head movements in light, could account for the observed phenomena. The two main classes of inhibitory neurons targeting RS cells in V1 are the parvalbumin (PV) and somatostatin (SOM) expressing cells^18^.

As described above (Supplementary Fig. 2) a fraction of FS cells (putative PV expressing cells), mainly those located in deep layers, were excited by head movements in the dark, potentially accounting for the suppression of V1 neurons across cortical layers^19^. However, those same FS cells were equally excited by head movements in light (Dark vs Light: vMI: 0.31±0.03 vs 0.25±0.04, n = 55 cells, P = 0.1, Supplementary Fig. 3c). Thus, PV cells are unlikely to mediate that component of V1 suppression to head movements that is relieved by light because they were equally excited by head movements in light and dark. We thus turned to SOM cells.

To isolate the response of V1 SOM cells to head movements, we opto-tagged these neurons via the conditional expression of Channelrhodopsin 2 in SOM-Cre mice^20^. Like RS and FS cells, SOM cells in the superficial layers were suppressed, on average, by head movements in the dark and excited by head movements in light (Fig. 3b,c). Therefore, like FS cells, superficial SOM cells are unlikely candidates for the head movement mediated suppression of V1 neurons in the dark and their facilitation by light. In striking contrast to FS cells and to SOM cells in superficial layers, SOM cells in deep layers were excited by head movements in dark and suppressed by the same head movements in light (Fig. 3b,c). Thus, unlike RS and FS cells, light did not facilitate their response to head movements but instead decreased their vMI. Furthermore, in response to light, i.e. prior to table rotations, the basal activity of these SOM cells increased, again unlike RS and FS cells. In fact, the larger the increase in their basal activity by light, the larger their decrease in vMI (Fig. 3d-f). These results show that SOM cells in deep layers are excited by light as well as by head movements in the dark, and that light occludes their excitation to head movements.

**Figure 3.**
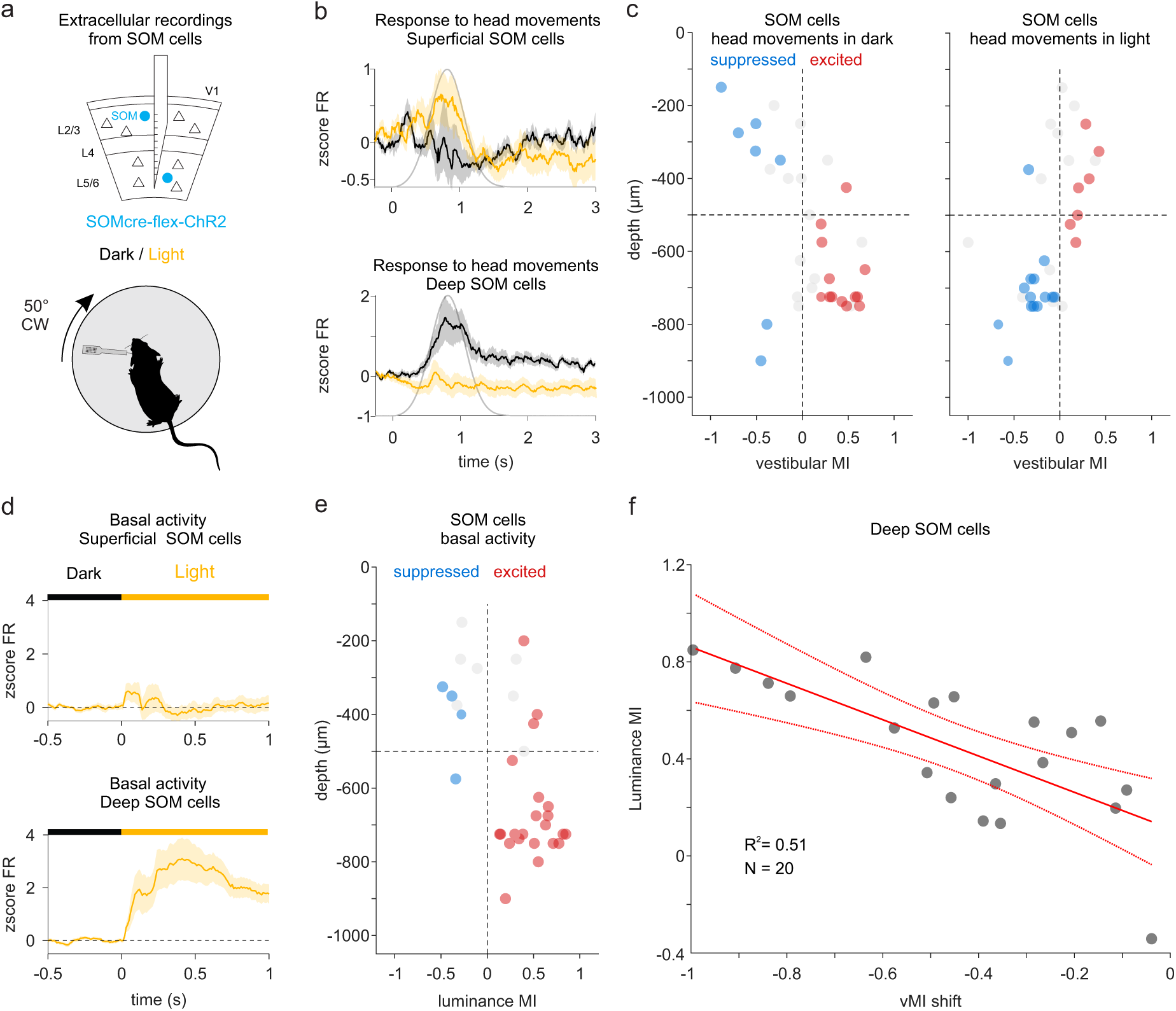
Excitation of layer 5 SOM cells by head movements in dark and suppression in light. **(a)** Experimental configuration. Top: Schematic of extracellular recordings from photo-tagged SOM cells across layers in V1. Bottom: The extracellular linear probe is inserted in the left V1 of the head-fixed, awake mouse to record the activity of SOM cells in response to clockwise (CW) rotation of the table, in dark or light. **(b)** Averaged z-score of the firing rate (zscore FR) of SOM cells during CW rotations in dark (black traces) and in light (yellow traces) for SOM cells recorded in the superficial layers (top panel; n = 12) and deep layers (bottom panel; n = 20). Shading is SEM. Gray trace is velocity profile. Note the opposite response in dark and light for SOM cells in superficial and deep layers. **(c)** Vestibular modulation index (vMI) to CW rotations of individual SOM cells plotted against cortical depth. Blue, red and gray circles are suppressed, excited and non-significantly modulated SOM cells, respectively. Left: vMI of SOM cells in dark; Right, vMI of the same SOM cells in light (Dark vs Light : Superficial layers vMI: −0.24±0.11 vs 0.10±0.07; P = 0.021; Deep layers vMI: 0.25±0.07 vs −0.23±0.06; P = 6.1e-05; n= 32 SOM cells; 15 recordings from 8 mice). Horizontal dotted line indicates approximate border between superficial and deep layers (between layer 4 and 5). Note shift of deep layer SOM cells towards negative vMIs. **(d)** Averaged z-score of the firing rate (zscore FR) of SOM cells in response to light onset for SOM cells recorded in the superficial layers (top panel; n = 12) and deep layers (bottom panel; n = 20). Shading is SEM. Horizontal bar indicates time of dark/light transition (time 0). Note the strong increase in firing rate of deep layer SOM cells by light. **(e)** Luminance modulation index of SOM cells plotted against cortical depth. Blue, red and gray circles represent SOM cells whose basal activity is suppressed, excited and non-significantly modulated by light onset, respectively (Superficial layers lumMI: −0.009±0.11; Deep layers lumMI: 0.44±0.06, same cells as in C). Horizontal dotted line indicates approximate border between superficial and deep layers. **(f)** vMI shift (vMI in light - vMI in dark) plotted against lumMI. Continuous red line: linear regression (R^2^ = 0.51; n = 20). Dotted red lines: confidence interval of the linear regression. Note negative values of vMI shift as compared to RS cells (Fig. 2g) yet similar inverse relationship between vMI shift and lumMI.

This observation makes SOM cells in deep layers potential candidates for the suppression of V1 neurons by head movement in dark, because these SOM cells are excited by head movements in dark, and for the facilitation by head movement in light, because these SOM cells are no longer excited by head movements in light. This result also makes deep-layer SOM cells potential candidates for the suppression of the basal activity of V1 neurons by light.

### SOM cells suppress V1 neurons in response to head movements in dark but not in light

To directly test the involvement of SOM cells in the head movement mediated response of V1 neurons, we specifically ablated these cells in V1 through the conditional expression of Caspase-3 in SOM-Cre x Ai14 mice. Histological analysis demonstrated an 86.5±0.02% reduction of SOM cells compared to the contralateral non-injected hemisphere (Fig. 4b, see methods). SOM cell ablation slightly increased the basal activity of RS cells (control average FR = 2.87±0.16Hz, SOM ablation FR = 3.30±0.17Hz, P = 0.0004).

**Figure 4.**
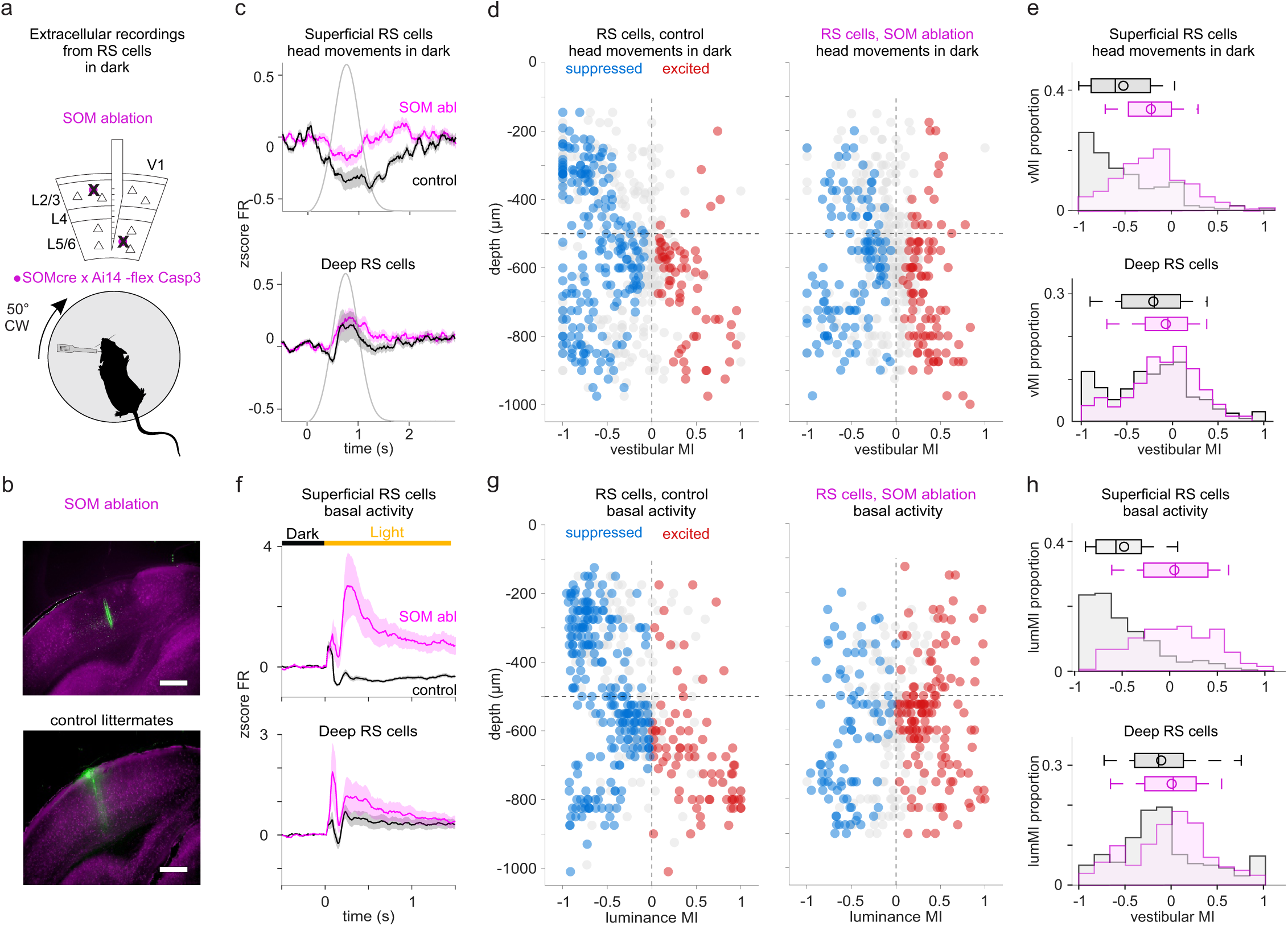
SOM cells suppress RS cells in response to head movements and light onset. **(a)** Experimental configuration: Top: Schematic of extracellular recordings from V1 in which SOM cells have been ablated. Bottom: The extracellular linear probe is inserted in the left V1 of the head-fixed, awake mouse to record the activity of RS cells in response to clockwise (CW) rotation of the table in dark. **(b)** Fluorescence microscopy images of V1 coronal sections (SOM Cre x Ai14 mouse) with (top panel) or without (bottom panel) the conditional expression of virally injected caspase-3. The electrode track is in green. Note the reduction in fluorescence around the recording site in the injected as compared to the un-injected hemisphere. Scale bar: 400 μm. **(c)** Averaged z-score of the firing rate (zscore FR) of RS cells during CW rotations in dark for SOM ablated animals (magenta) and un-injected, control littermates (black). Top panel: All RS cells recorded in superficial layers (n = 149 cells (magenta) and 182 cells (black)). Bottom panel: All RS cells recorded in deep layers (n = 339 cells (magenta) and 333 cells (black)). Shading is SEM. Gray trace is velocity profile. Note the reduction of the vestibular suppression in superficial RS cells and the increase in excitation in deep RS cells in SOM ablated animals as compared to un-injected littermates. **(d)** Vestibular modulation index (vMI) for CW rotations of individual RS cells plotted against cortical depth. Blue, red and gray circles are suppressed, excited and non-significantly modulated RS cells, respectively. Left: vMI of RS cells in non-injected littermates. Right: vMI of RS cells in SOM ablated animals (Control [n=11 mice] vs Ablation [n=9 mice], Superficial layers vMI: −0.56±0.03 [n = 182 cells] vs −0.20±0.03 [n = 149 cells], P = 3.18e-16; suppressed: 38±3% vs 25±4%, P = 0.027; excited: 4.8±1.4% vs 19±3%, P = 0.0023; Deep layers vMI: −0.22±0.025 [n = 333 cells] vs −0.073±0.02 [n = 339 cells], P = 3.1e-06; suppressed: 32±2% vs 21±1%, P = 0.001; excited: 19±3% vs 27±2%, P = 0.044; 1 recording per animal for both conditions). Horizontal dotted line indicates approximate border between superficial and deep layers (between layer 4 and 5). Note shift towards positive vMI in SOM ablated animals, especially in superficial layers. **(e)** Distribution of the vMI in SOM ablated animals (magenta) and control littermates (black) for RS cells recorded in superficial (top panel) and deep (bottom panel) layers. Box plots show median, first and third quartiles, minimum/maximum values and mean (circle). **(f)** Averaged z-score of the firing rate (zscore FR) of RS cells in response to light onset in SOM ablated animals (magenta) and control littermates (black). Top panel: All RS cells recorded in superficial layers (n = 99 cells (magenta) and 199 cells (black)). Bottom panel: All RS cells recorded in deep layers (n = 260 cells (magenta) and 276 cells (black)). Shading is SEM. Horizontal bar indicates time of dark/light transition (time 0). **(g)** Luminance modulation index (lumMI) of individual RS cells plotted against cortical depth. Blue, red and gray circles are suppressed, excited and non-significantly modulated RS cells, respectively. Left: lumMI of RS cells in un-injected littermates; Right: vMI of SOM ablated animals (Control [n=8 mice] vs SOM ablation [n=8 mice]: Superficial layers lumMI: −0.50±0.03 [n = 199 cells] vs 0.04±0.05 [n = 99 cells]; P = 8.5e-20; suppressed: 60±10% vs 25±6%, P = 0.015; excited: 4.6±1.6% vs 41±10%, P = 0.005; Deep layers lumMI: −0.09±0.03 [n = 270 cells] vs −0.01±0.03 [n = 260 cells], P = 0.003; suppressed: 42±8% vs 27±4%, P = 0.05; excited: 15±4 vs 41±5%, P = 0.002, 1 recording per animal). Horizontal dotted line indicates approximate border between superficial and deep layers. Note shift towards positive lumMI in SOM ablated animals. **(h)** Distribution of the lumMI in SOM ablated animals (magenta) and control littermates (black) for RS recorded in superficial (top panel) and deep (bottom panel) layers. Box plots show median, first and third quartiles, minimum/maximum values and mean (circle).

Ablation of SOM cells significantly reduced the head movement mediated suppression of V1 neurons in the dark, reversed the suppressive impact of light on basal activity, and reduced the facilitation of the V1 response to head movements in light (Fig. 4). The reduction of head movement mediated suppression in dark was particularly prominent for RS and FS cells in superficial layers (Fig. 4 and Supplementary Fig. 5b-d for FS cells), consistent with our observation that neurons in superficial layers are more suppressed by head movements than those in deep layers (Fig. 1 and Supplementary Fig. 2). SOM cell ablation also profoundly impacted the light-mediated suppression of V1 basal activity (Fig. 4f-h). In Caspase-3 injected animals, light increased instead of suppressing basal activity (Fig. 4f-h and Supplementary Fig. 5e-g for FS cells). If the relationship between the light-mediated suppression of basal activity and the facilitation of the response to head movements is causal, reducing the impact of light on basal firing rate should decrease the facilitation of the response to head movements by light. Indeed, facilitation of responses to head movements in light was strongly reduced in SOM cell ablated animals (Fig. 5a-d), as compared to their control littermates (Fig. 5e-h).

**Figure 5.**
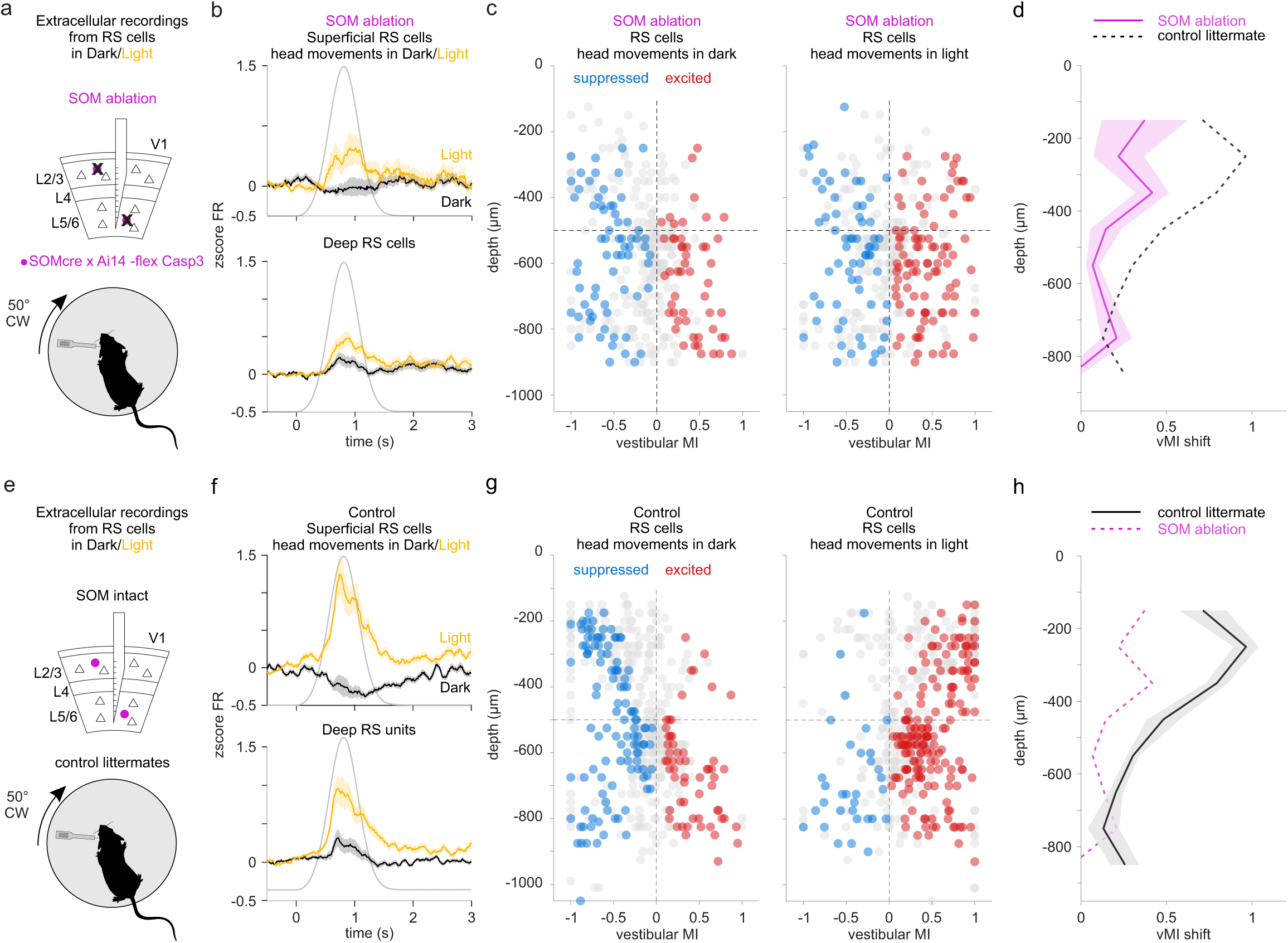
SOM cells contribute to the light-mediated facilitation of the response to head movements. **(a)** Experimental configuration: Top: Schematic of extracellular recordings from V1 in which SOM cells have been ablated. Bottom: The extracellular linear probe is inserted in the left V1 of the head-fixed, awake mouse to record the activity of RS cells in SOM ablated mice in response to clockwise (CW) rotation of the table in dark and light. **(b)** Averaged z-score of the firing rate (zscore FR) of RS cells recorded in SOM ablated mice during CW rotations in dark (black) and light (yellow). Top panel: All RS units recorded in superficial layers (n = 99 cells, magenta). Bottom panel: All RS cells recorded in deep layers (n = 260 cells, black). Shading is SEM. Gray trace is velocity profile. **(c)** Vestibular modulation index (vMI) for CW rotations of individual RS cells recorded in SOM ablated mice plotted against cortical depth. Blue, red and gray circles are suppressed, excited and non-significantly modulated RS cells, respectively. vMI of RS cells in dark (left) and light (right; n = 359 RS cells; 8 mice; 1 recording per mouse). Horizontal dotted line indicates approximate border between superficial and deep layers (between layer 4 and 5). Note shift towards positive vMI in SOM ablated mice. **(d)** vMI shift (vMI in light - vMI in dark) plotted against cortical depth. Magenta: SOM ablated mice; Black dotted line: control littermates (see (h)). Shading is SEM. (Control [n=8 mice] vs SOM ablation [n=8 mice]: vMI shift in superficial layers: 0.79±0.04 [n = 199 cells] vs 0.26±0.06 [n = 99 cells], P = 7.2e-13; vMI shift in deep layers: 0.25±0.03 [n = 276 cells] vs 0.08±0.03 [n = 260 cells], P = 8e-07). Note the reduction in vMI shift in SOM ablated animals. **(e-f)** as in (a-d) but for control non-injected littermates with intact SOM cells (n = 475 RS cells (199 cells in superficial layers and 276 cells in deep layers; 8 mice; 1 recording per mouse). The black line in (F) is the vMI shift in control littermates and the magenta dotted line the vMI shift in SOM ablated mice (from (D)).

Taken together, these data show that the suppression of V1 neurons by head movements in dark and the dependence of this suppression on ambient luminance relies on SOM cells: SOM cells suppress V1 neurons in response to head movements in dark and, by releasing their suppression in light, enable facilitation of the response. Furthermore, these data show that SOM cells also suppress V1 basal activity in response to light, thus explaining the observed relationship between the magnitude of suppression of basal activity by light and the degree of facilitation of the response to head movements (RS cells: Fig. 2g and FS cells: Supplementary Fig. 3g).

### PV and 5HT3aR cells do not contribute to the suppression of V1 by head movements

We verified that the impact of SOM cell ablation on head movement and light responses in V1 was not due to a nonspecific decrease in inhibition, by selectively ablating PV cells and 5-Hydroxytryptamine3a receptor (5HT3aR) expressing inhibitory cells, which together with SOM cells account for nearly 100% of inhibitory neurons in cortex^21^. We confirmed the ablation of PV cells both histologically (92±0.04% ablated PV cells) and by the reduction of recorded FS cells (11±2.2% vs 2.9±1.2%, P = 0.03; n = 5 mice for both). Ablation of PV cells led to a small increase in basal activity as compared to non-injected areas in V1 (2.46±0.29 Hz vs 2.91±0.14 Hz, P = 0.003). Ablation of 5HT3aR cells reduced this neuronal population by 86.2±0.008% without any significant impact on basal activity of RS cells (2.80±0.16Hz vs 2.69±0.14Hz, control: n = 8 mice; 5HT3aR cell ablation: n = 11 mice).

Neither PV nor 5HT3aR cell ablation significantly affected the response of RS cells to head movements (Supplementary Fig. 6a-d and Supplementary Fig. 6e-h, respectively). Moreover, neither PV nor 5HT3aR cell ablation impacted the suppression of basal activity by light (PV control: lumMI = −0.21±0.03; n = 272 cells; 5 mice; PV ablation: lumMI = −0.16±0.03; n = 253 cells; 4 mice; p = 0.17; 5HT3aR control: lumMI = −0.22±0.02; n = 631 cells; 10 mice; 5HT3aR ablation: lumMI = −0.18±0.02; n = 583 cells; 10 mice; p = 0.22). Therefore, PV and 5HT3aR cells do not significantly contribute to either the head-movements mediated suppression of RS cells or to the light mediated suppression of their basal activity. These results indicate that the impact of SOM cell ablation on V1 response to head movements and to light is not due to a non-specific decrease in inhibition. Instead, these results highlight the unique role of SOM cells in the suppression of V1 by head movements in dark, the relief of suppression to head movements in light, and the suppression of V1 basal activity by light.

### Light mediated facilitation of V1 activity to head movements in freely moving mice

Although head-fixed mice on a rotating table allow us to stimulate the vestibular organs through controlled head movements, in freely moving animals the vestibular signal arise from head movements initiated by the animal. To determine whether these active head movements impact V1 activity in a similar manner to that observed in response to head movements triggered by passive table rotations, we performed electrophysiological recordings from V1 in freely moving animals. We recorded the angular velocity of head movements using an inertial measurement unit attached to the head of the animal (Fig. 6a, see methods). As above, the eyelids were sutured to prevent patterned visual stimulation, and dark/light conditions were alternated every two minutes. Spontaneous exploratory behavior was accompanied by a wide range of head angular velocities along the three orthogonal planes captured by the IMU, with higher velocities for the horizontal component of the head movements (yaw) (Fig. 6b). The distribution of angular velocities was similar in dark and light conditions (yaw average head angular velocity in dark vs. light: 46.7±4.0 °/s vs 46.8±5.5 °/s, P = 1, n = 4 mice). To characterize the relationship of firing rate to angular velocity, we calculated the head rotation modulation index (hrMI) for yaw rotations (see methods). A large fraction of V1 cells (53%) was significantly modulated by angular velocity in the dark, around any plane and direction and 49% of cells specifically by head motions with a yaw component (CW and CCW). Consistent with head fixed conditions, average V1 activity was suppressed by active head movements in the dark, with a more robust suppression of cells in superficial layers (Fig. 6c,d). Furthermore, head movements in light resulted in an overall shift towards facilitatory responses leading to and a 25±8% decrease in suppressed cells and a 33.3±11.8% increase in facilitated cells (Fig. 6e).

**Figure 6.**
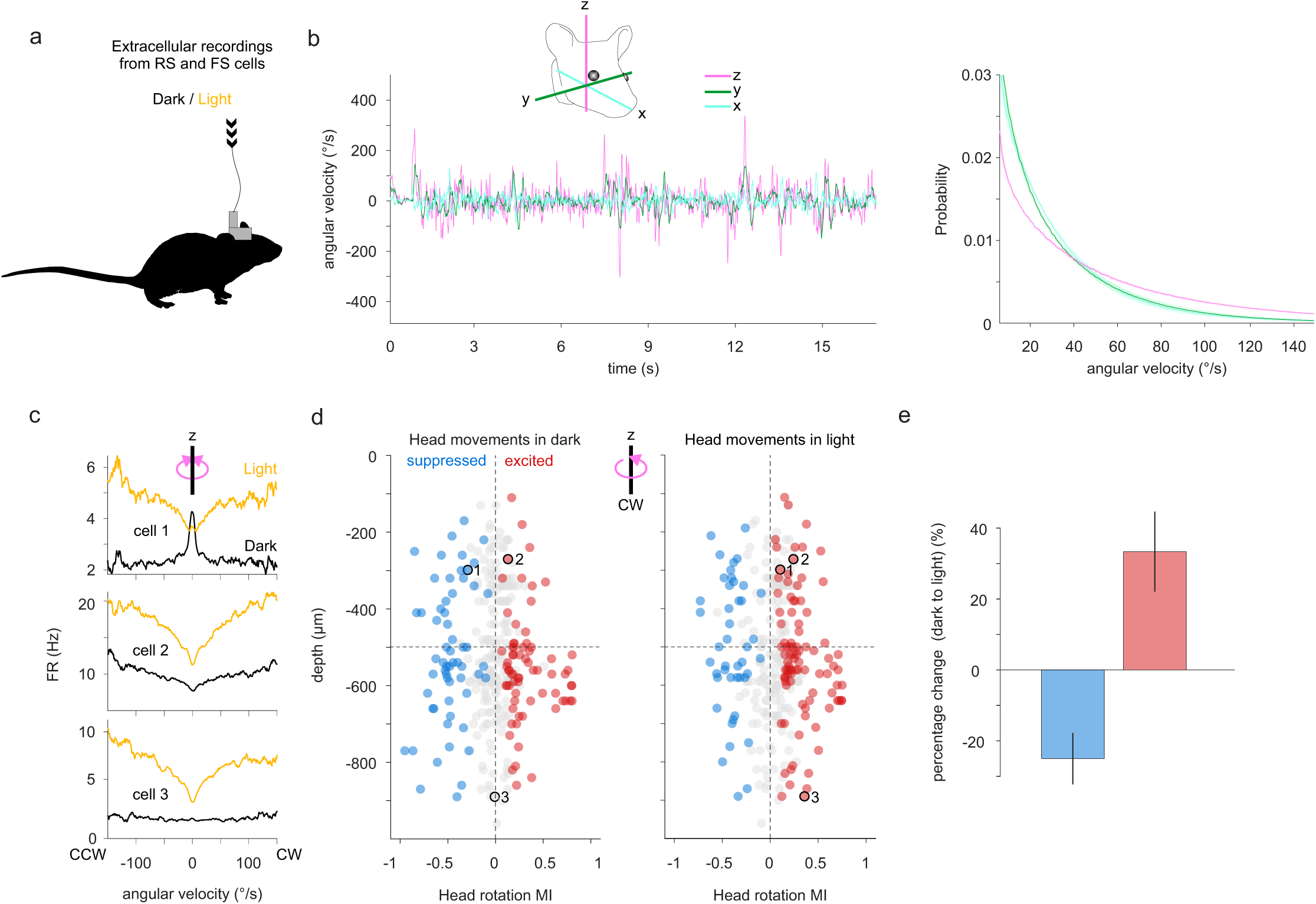
Light mediated facilitation of V1 activity to head movements in freely moving mice. **(a)** Experimental design: A chronic extracellular linear probe is inserted in the left primary visual cortex (V1) of a freely moving mouse. Head motion is monitored with an IMU system attached to the head of the animal. Light and dark condition are alternated every 2 minutes. The light source homogeneously illuminates the arena (see Methods). **(b)** Left: Example traces illustrating head angular velocity in time around 3 orthogonal axes of rotation. Inset: Schematic illustration of the three axes relative to the head of the mouse. Right: Probability distribution of angular velocity around the three axes of rotation. Shading is SEM. **(c)** Firing rate (FR) plotted against angular velocity around the z-axis (yaw) for 3 example cells in dark (black traces) and in light (yellow traces). Positive and negative values are clockwise (CW) and counterclockwise (CCW) rotations, respectively. Note that Cell 1 is suppressed by yaw in dark and excited by yaw in light, Cell 2 is more excited by yaw in light than in dark and Cell 3 is does not respond to yaw in dark but is excited by yaw in light. **(d)** Head rotation modulation index (hrMI) for CW rotations of individual isolated cells plotted against cortical depth. Blue, red and gray circles are suppressed, excited and non-significantly modulated units, respectively. Left: hrMI of isolated cells in dark; Left, hrMI of the same cells in light (n = 324 cells; 4 recordings from 4 mice). Black circles illustrate examples units from (c). Horizontal dotted line indicates 500 μm depth. Note shift towards positive hrMI in light. Cortical depth is measured from the pia. **(e)** Change in the percentage of suppressed (blue) and excited (red) cells in response to CW head rotations around the z-axis from dark to light. Note the decrease in suppressed (−25±8%) and the increase in exited units (33.3±11.8%). Percentages are mean ± SEM.

These results show that active head movements control the activity of large fractions of V1 neurons in a luminance dependent manner. On one hand, active head movements in the dark on average suppress V1 neurons, similar to passive head movements triggered by table rotations. Active head movement in light, on the other hand, on average excites V1 neurons, again similar to passive head movements.

## Discussion

Movements of an animal through its environment continuously activate the animal’s vestibular system whose signal are broadcast throughout the brain, including visual cortex. These signals are believed to contribute to the organism’s ability to distinguish between sensory stimuli resulting from the animal’s movement in the environment from those resulting from an actual change in the environment. This study shows that head movements control the activity of a majority of V1 neurons throughout cortical layers and that this control occurs in a luminance dependent manner. While head movements in the dark mainly suppress V1 neurons, the same head movements in light excite V1 neurons. The suppression of V1 neurons in the dark is largely mediated by SOM cells which are excited by head movements. In light, however, the reduced excitation of SOM cells by head movements relieves the suppression of V1 neurons. Thus, this study reveals a light-dependent switch in the response of V1 to head movements and identifies a circuit in which SOM cells are key integrators of vestibular and luminance signals.

The strong suppression of V1 by head movements in darkness may act as a veto in the processing of visual stimuli, for example the shift of the visual scene as the animal moves its head, that are unreliable due to dim ambient luminance conditions. In contrast, excitation of V1 by head movement in light may contribute to the combination of visual and vestibular signals as the animal navigates through its environment in bright conditions, enabling visual cortex to attribute shifts in the visual scene to head movements of the animal.

Head movements similarly impacted both RS and FS cells across layers, putative pyramidal and PV cells, respectively. Both cell types were overall suppressed in dark and excited in light by head movements. Furthermore, the impact of light on V1 basal activity was also similar between RS and FS cells, being predominantly suppressive, especially in superficial layers. Interestingly, both RS and FS cells are targeted by SOM cells^18^, making the latter cell-type a potential candidate for both the head-movements and the light mediated suppression of V1 activity. Consistent with this possibility, deep-layer SOM cells increased their firing rate in response to head movements in dark and in response to light onset, as would be expected for neurons suppressing their targets in response to these two distinct stimuli. Further supporting this possibility, SOM cell ablation reduced both the head-movement and the light mediated suppression of basal activity in RS and FS cells. Thus, our data show that SOM cells contribute to the head movement mediated suppression of V1 neurons in the dark. Our data also demonstrate that SOM cells represent the basis for the light mediated suppression of basal activity in V1^16, 17^.

Importantly, RS, FS and SOM cells all showed an inverse relationship between the impact of light on basal firing rate and head-movements mediated responses: For RS and FS cells, decreasing their basal firing rate with light facilitated their response to head movements (Fig. 2). For SOM cells, increasing their basal firing rate with light reduced their excitation by head movements (Fig. 3). Thus, while the impact of light is opposite between RS and FS cells on one hand and SOM cells on the other, the relationship between basal firing rate and responses to head movements is similar. If this relationship is causal, reducing the impact of light on basal firing rate should decrease the facilitation of the response to head movements by light. This is exactly what we observed upon ablating SOM cells: We not only removed the light-mediated suppression of RS and FS basal firing rate, but also reduced the facilitation of their response to head movements by light. Thus, SOM cells represent the causal link between the light mediated decrease in basal firing rate and the correlated facilitation of responses to head movements observed in V1 neurons.

Although SOM cell ablation experiments encompassed all cortical layers, our data suggest that SOM cells located in deep layers are those that mainly contribute to the suppression of V1 in response to head movements in dark, to the relief of suppression by light, and to the reduction in basal activity by light. In fact, by comparing the activity of SOM cells across layers, only SOM cells in deep layers were excited in response to head movements in the dark, no longer so in light, and increased their basal activity in light. These SOM cells are likely deep layer Martinotti cells, because of their location and spike shape^22–24^ (see methods). Therefore, we propose that deep layer Martinotti cells integrate vestibular and luminance signals in V1.

Future work will determine the source of vestibular input onto SOM cells. We show that the impact of head movements on V1 activity depends on the vestibular organ, since its surgical ablation abolished the head-movement mediated response in V1. It has been suggested that retrosplenial cortex is a source of vestibular input to V1^4^. However, several other areas responding to vestibular stimulation have been shown to project to V1^3, 25, 26^. Also, the nature of the luminance input onto SOM cells remains to be established. While a large fraction of neurons in the dorsolateral geniculate nucleus of the thalamus (dLGN), the primary visual input to V1, increases firing in response to increases in luminance^16, 27^, SOM cells are not a major target for dLGN afferents^28^. However, a fraction of local layer 5 RS neurons increase their basal firing rate with light, and since these neurons excite deep layer SOM cells^29^, they may represent the SOM cells’ luminance input.

The impact of SOM cell ablation on V1 responses to head movements was not due to a general decrease in inhibition. Ablation of the two other major groups of cortical inhibitory neurons, the PV and 5HT3aR cells, did not impair the suppression of V1 by head movements in the dark, the facilitation by light or the effect of light on basal activity. This points to a very specific role of SOM cells as integrators of head-movements and luminance. Interestingly, other non-visual modulations of V1 activity have been shown to be affected by light, like the response of V1 to auditory stimuli^12^. Future work will determine whether SOM cells represent a general mechanism for the luminance dependence of non-visual responses in V1.

Despite the overall suppression of V1 in response to head movements in the dark and the overall excitation in light, the magnitude of suppression or excitation varied across individual neurons. This variability may reflect the different tuning properties of individual cells for different axes of head motion, a possibility that will be tested by moving the animal along different planes. This variability may also correlate with the tuning properties of individual neurons to visual stimuli, a possibility that can be determined by comparing the response of individual neurons to head movements with their visual tuning properties.

In conclusion, our work reveals that head motion exerts a strong control on the activity of V1 neurons in a layer and cell-type specific manner and that ambient luminance controls the sign and magnitude of this impact through SOM cells. The impact of the vestibular system on V1 and its luminance dependence suggest that the landscape of activity in V1 changes continuously with the temporal dynamics of the animal’s motion through its environment. An ethological understanding of how V1 processes visual information will, thus, necessitate a thorough elucidation how these head movement generated moment-to-moment fluctuations in V1 activity are integrated with the ongoing flow of visual signals.

## Acknowledgments

We thank all the members of the Scanziani and the Nelson lab for discussions about the project and comments on the manuscript; A. Nelson, R. Nicoll, B. Barbour, S. Dieudonné, N. Rebola and B. Liu for critical reading of the manuscript; and M. Mukundan, J. Lee, B. Wong, L. Bao, Y. Li, and O. Lahrach for technical support.

## Contributions

G.B. and M.S. designed experiments and wrote the manuscript. G.B. performed all the experiments, except Y.S. implanted the chronic electrodes. G.B. analyzed the data.

## Methods

All experimental procedures were conducted in accordance with the regulations of the Institutional Animal Care and Use Committee of the University of California, San Francisco.

### Animals

Data were collected from male or female C57BL/6J mice or from heterozygous SOM-Cre x Ai32 (Som-Cre: JAX#028864; Ai32: JAX**#**024109), SOM-Cre x Ai14 (Som-Cre: JAX#028864; Ai14: JAX**#**007914), PV-Cre x Ai14 (PV-Cre: JAX#017320; Ai14: JAX**#**007914), 5HT3a-cre x Ai14 (5HT3a-Cre: MMRRC#036680-UCD; Ai14: JAX**#**007914) transgenic animals. Mice were housed on a reverse light cycle. All animals were older than 2 months at the start of experiments.

### Viruses

The following adeno-associated viruses (AAV) were used: AAV1-EF/a-Dio-Casp3-2A-TEV (final titer: 2.1.10^12^ GC mL^-1^) and AAV1.Ef1α.DIO.ChR2(H134-R).eYFP.WPRE.hGH (final titer: 4.10^12^ GC mL^-1^), Univ. of Pennsylvania Viral Vector Core.

### Surgical procedures

#### Viral Injections

Mice were anesthetized with 2% isoflurane and placed in a stereotactic apparatus (Kopf). Core body temperature was maintained at 37°C with a rectal probe and a heating pad (FHC). A thin layer of lubricant ointment (Rugby Laboratories) was applied to the eye, the head was shaved and disinfected with povidone iodine and 2% lidocaine solution was administered subcutaneously at the incision site. A craniotomy (approx. 300 µm in diameter) was performed with a micro-burr (Gesswein) mounted on a dental drill (Foredom). Viral suspensions were loaded in beveled glass capillaries (tip diameter: 15-30 µm) and injected with a micropump (UMP-3, WPI) at a rate of 20-30 nl/minute into the parenchyma. The pipette was removed from the brain 10 minutes after the completion of the injection, the head plate was attached just after the virus injection and 0.1 mg/kg buprenorphine was administered subcutaneously as a postoperative analgesic.

This virus was injected in the left V1 at 3 sites forming a triangle (2.3 mm medio-lateral/0.45 mm anterior to the lambdoid suture, 2.8 mm medio-lateral/0.45 mm anterior to the lambdoid suture, 2.55 mm medio-lateral/0.8 mm anterior to the lambdoid suture) at 2 depths for every injection site (250 and 500 µm). Experiments were performed at least 3 weeks after the virus injection. For PV cell ablations (AAV1-EF/a-Dio-Casp3-2A-TEV in PV Cre x Ai14 mice), we injected only one site and two depth (250 and 500 µm) to reduce the extent of PV cells ablation and hence minimize the risk of seizures.

#### Eyelid Suturing

Mice were anesthetized with 2% isoflurane. The eye was flushed and eyelids swabbed with dilute betadine ophthalmic solution. Proparacaine ophthalmic solution was applied to numb the eye. The eyelid margins were then sutured closed with 1 mattress suture using either polypropylene or vicryl suture.

#### Head Plate Implantation for Head-fixed Recordings

Mice were implanted with a T-shaped head-bar at least 2.5 weeks before the day of the recording. The animals were anesthetized with 2% isoflurane, the scalp was removed and the skull was disinfected with alcohol and povidone iodine and scored with bone scraper. The edge of the skin was glued to the skull and the metal head-bar was sterilized and mounted using dental cement (Ortho-Jet powder; Lang Dental) mixed with black paint (iron oxide) or Relyx Unicem2 automix (3M ESPE). The head-bar was stereotactically mounted with the help of an inclinometer (Digi-Key electronics 551-1002-1-ND). The inclinometer was instrumental in calibrating the angle of the two axes of the T-shaped head bar in relation to the sagittal and medio-lateral axes of the head. Following the bar implantation, black dental cement was used to build a recording well surrounding the recording site. The surface of the skull above the left visual cortex was not covered with dental cement, but was coated with a thin layer of transparent cyanoacrylate glue. Animals were injected subcutaneously with 0.1 mg/kg buprenorphine and checked daily after the head-bar surgery. For at least 5 days before recording, mice were habituated to head fixation within the recording setup.

#### Craniotomy for Electrophysiological Recordings

On the day before recording, mice were anesthetized with 2% isoflurane and the skull above the recording sites was drilled off. The dura was not removed, and the exposed brain was kept moist with artificial cerebrospinal fluid (ACSF; 140mM NaCl, 5mM KCl, 10mM D-glucose, 10mM HEPES, 2mM CaCl2, 2mM MgSO4, pH 7.4). V1 recordings were performed around 2.6 mm medio-lateral/0.6 mm anterior to the lambdoid suture.

#### Chronic Electrode Implantation

Mice were implanted with recording electrodes under 2% isoflurane anesthesia. These procedures were performed in two steps. First, a head plate base was placed around the implantation target area. After 7 days of recovery, mice were habituated at least 5 days to freely explore the arena (see below) with the head attached to the cables for the inertial measurement unit (IMU) and the electrophysiological recording. We also added approximately 3g weight on the head to mimic the presence of the microdrive, the IMU and the electrode. After this period of habituation to the setup, mice underwent the second step, namely the silicon probe implantation procedure. Just before performing the chronic electrode implantation, we stitched the eyelids of both eyes (see above). A single- or two-shank silicon probe (Cambridge NeuroTech H3 64×1 or Diagnostic Biochips P128-3, respectively) was mounted on a movable microdrive to record V1 neuronal activity. After the ground electrode implantation (0.005” diameter stainless wire, A-M systems) in the cerebellum and the craniotomy above the target implantation site, the probe was implanted at the same stereotaxic coordinate as the acute recording. The probe was lowered to 0.8-0.9 mm below the brain’s surface. After the recovery from the isoflurane anesthesia, the probe depth was finely adjusted by moving the microdrive so that the recording channels could cover the entire depth of V1. The experiments were performed the day after implantation.

#### Bilateral Vestibular Lesions

For bilateral lesion experiments, mice were anesthetized with 2% isoflurane. Core body temperature was maintained at 37°C with a rectal probe and a heating pad (FHC). The lateral regions of the posterior and horizontal semicircular canals were exposed. The canal bone was then thinned until punctured and a microfiber needle inserted to deliver kanamycin (5 μl; 50 mg/g b.w). The skin was sutured with a few stitches of 6/0 suture. Mice recovered for three to five days in their home cages before recording of V1 activity.

### Electrophysiology

Extracellular recordings were performed using the following silicon probes Neuronexus:A1×32-5mm-25-177-A32;A1×32-Edge-5mm-20-177-A32; A2×32-5mm-25-177-A64, 1×64-Poly2-6mm-23s-160 or Cambridge Neurotech: ASSY-77 H2 (Acute 64 channel H2 probe, 2 shanks @250um, 8mm length), ASSY-77 H5 (Acute 64 channel H5 probe, 1 shank, 9 mm length). The recording electrodes were controlled with Luigs & Neumann micromanipulators mounted on the rotating platform (see below) and stained with DiI or DiO lipophilic dyes (Life Technologies) for post hoc identification of the electrode track. We recorded the signals at 30 kHz (head fixed recordings) or 20 kHz (recordings in freely moving animals) using an INTAN system (RHD2000 USB Interface Board, INTAN Technologies).

### Head-fixed rotations

The animal was head-fixed with a ∼ 20 degrees angle along the naso-occipital axis of the head, to place the horizontal vestibular canal approximately parallel to the rotating platform. The platform was attached to a gearbox 15:1 (VTR010-015-RM-71 VTR, Thomson) that increased the torque of a servo motor (AKM53L-ANC2C-00 KEC0432 AC Servomotor 1.83kW, Kolmorgen). The motor was tuned using a servo drive (AKD-B013206-NBAN-0000 servo drive, Kolmorgen) and controlled in velocity mode using analog waveforms computed in Labview. To control the velocity and amplitude of head movements, we fixed the head of awake mice in the center of a servo-controlled table enabling the rotation of the animal along the horizontal plane (50 degrees rotation; 80°/s peak velocity; unless stated otherwise Fig. 1a).

### Illumination

To illuminate homogeneously the right eye, we placed a light diffuser (DG20-600, Thorlabs) 3-5 cm away from the eye, and an optical fiber (core = 960µm / NA = 0.63) coupled to a blue LED (470 nm; Doric Lenses) was placed 6-7 cm away from the diffuser. The light source, as well as the light diffuser, were attached to the rotating platform, to move together with the animal. The illumination power at this distance was between 0.75 - 1 mW, the light was immediately turn on at this power and turned off by ramping down the power over 0.5 s to limit rebound activation. Illumination was randomized and lasted 4 seconds. During a typical trial the rotation velocity peaked 2 seconds after the onset of the illumination. As the illumination was randomized, the interval between 2 illuminations was variable with a minimum of 10 s.

### SOM cells Photo-tagging

ChR2-expressing SOM were activated via an optical fiber (core = 960µm/ NA = 0.63) coupled to a blue LED (470nm; Thorlabs) and placed above V1. Light from the fiber optic spanned the entire area of V1 (Power: 5-6 mW measured at the fiber tip). LED power was measured before each experiment with a digital power meter (PM100D, Thorlabs). The photo-tagging of the cells was performed before and after vestibular stimulation protocol.

### Arena for freely moving animals

The animal navigated inside a circular arena (LED light Sheet panel, Tap plastic; dimension 24 cm diameter/ 32 cm height) with an inertial measurement unit (IMU) placed parallel to Bregma-Lambda axis. Electrodes were connected to an assisted electrical rotary joint commutator (AERJ_12_HARW, Doric) with a RHD2000 0.9m ultra-thin SPI cable (INTAN, Part #C3213) using an omnetics adapter (ADAPTER_HO12, Doric). The IMU cable passed through the motorized commutator and connected to a non-motorized commutator (Slip ring Flange - 22mm diameter, Adafruit) that was placed on top of the motorized one. As both cables were attached together, the motorized commutator triggered the rotation of both cables together to compensate for head motion. The inertial signal was sent to a DAQ card and data was acquired using a Labview software, whereas electrophysiological signal was sent to the INTAN board. Light (∼70 cd/m2) and dark conditions were obtained using a flexible LED strip (Ustelar) attached to a larger arena (40 cm diameter) around the behavioral arena to obtain a homogeneous illumination within the behavioral arena. Sequences of 2 minutes illumination and darkness were alternated randomly, controlled via Labview.

### Tracking head movements in freely moving animals

To track the head angular velocity in freely moving mice, we attached an inertial measurement unit (IMU) that detected linear acceleration and angular velocity. The movement sensor board was built around the TDK/Invensense MPU-9×50 9 axis movement sensor^30^. The movement sensor USB interface supplied the sensor power, configuration, and data acquisition. Angular velocity data were acquired at 300Hz. The USB interface sent a synchronization input used to timestamp the electrophysiological recording by sending a signal to the INTAN board.

### Histology

For anatomical analysis, mice were transcardially perfused with phosphate buffered saline (PBS) and then with 4% paraformaldehyde (PFA) in PBS. For most of the post-mortem electrode tracks reconstructions, animals were not perfused. Brains were extracted from the skulls, post-fixed in 4% PFA overnight at 4°C and subsequently cut with a vibratome to 80-100 µm thick sequential coronal sections. Slices were collected and mounted in ProLong Gold (Life Technologies) or Vectashield mounting medium containing DAPI (Vector Laboratories H1500). Bright-field and fluorescence images were acquired using an Olympus MVX10 MacroView microscope. For quantification of the cell-specific ablation induced with caspase-3 virus injection in V1, cells were counted for every mouse over a distance of 800 μm mediolateral and 400 μm anteroposterior. This area was centered around the recording site and the number of cells in ablated areas was compared to the non-ablated right hemisphere from the same animals. For PV cell ablation, cells were counted for every mouse over a distance of 400 μm mediolateral and 400 μm anteroposterior.

### Data Analysis

#### Unit isolation

Automated spike sorting was then carried out using KiloSort (https://github.com/cortex-lab/Kilosort) by manual curation of the units using Phy (http://phy-contrib.readthedocs.io/en/latest/template-gui/). Single units were identified and all following analysis was carried out via Matlab (MathWorks). We excluded units with refractory period violations greater than 1%.

#### Response to platform rotations

The baseline spike rate was calculated on individual trials by averaging the spike rate over a window of 580 ms recorded when the platform was stationary. The spike rate in response to the rotation of the platform was calculated on the same trials by averaging the spike rate over a window of 580 ms centered around the peak of the rotation velocity profile. We pseudo-randomly alternated clockwise (CW; contraversive relative to the recording site) with counterclockwise (CCW; ipsiversive) rotations. Wilcoxon signed-rank tests were then applied to determine if a cell was significantly modulated (P<0.05) by the rotation of the platform. To compare the vMI collected from 2 populations of mice, we performed Wilcoxon rank sum tests. When reporting average vMIs, all cells were included, regardless of whether or not they were significantly modulated by the vestibular stimulus.

To compare the proportion of suppressed or excited cells, we computed the percentage of cells suppressed or excited for each animal in a given condition and performed a Wilcoxon rank sum test for both suppressed and excited conditions. Wilcoxon signed rank tests were performed instead to compare values obtained in the same recording (i.e. comparing dark and light conditions). Because both the fraction of modulated cells (Supplementary Fig. 1a) and the absolute vMI values were similar for CW and CCW rotations (median absolute vMI: 0.39±0.01 and 0.39±0.01 for CW and CCW rotations, respectively; P = 0.25, n = 928 cells), we focused on CW rotations exclusively.

#### Response to light

The spike rate before light onset was calculated on individual trials by averaging the spike rate over a time window of 500 ms before the onset of the light. The spike after light onset was calculated on the same trials by averaging the spike rate over a 500 ms window beginning 500 ms after light onset. Wilcoxon signed-rank tests were then applied to determine if a cell was significantly modulated (P<0.05) by light. To compare the lumMI collected from 2 populations of mice, we performed Wilcoxon rank sum tests. When reporting average lumMIs, all cells were included, regardless of whether or not they were significantly modulated by light.

#### Direction Preference Index

We define the direction preference index as (Rcw - Rccw)/(Rcw + Rccw), where Rcw and Rccw are the response of individual cells to clockwise and counterclockwise rotations (Supplementary Fig. 4d).

#### Definition of RS and FS cells

Cells were classified as fast or regular spiking based on properties of their average waveforms, at the electrode site with largest amplitude^31^. Briefly, three parameters were used for discrimination: the height of the positive peak relative to the initial negative trough, the time from the minimum of the initial trough to maximum of the following peak, and the slope of the waveform 0.5 ms after the initial trough. Two separable clusters were found, corresponding to fast-spiking (putative inhibitory) and regular-spiking (putative excitatory) neurons. These clusters were separated using a k-means clustering method. The same analysis was performed for both head fixed and freely moving recordings.

#### SOM cells analysis

Cells were considered to be putative SOM cells when their activity during the blue LED illumination of V1 triggered a significant increase of firing rate within the first 6 ms illumination (P<0.05, Wilcoxon signed rank test). For our analysis, we included only SOM cells with regular spike shape (see section: Definition of RS and FS cells). We excluded from our analysis five putative SOM cells that had fast spiking shape because they are non-Martinotti cells^22–24^ (see Discussion).

#### Head rotation modulation index in freely-moving mice

Spikes from isolated units were binned at the acquisition frequency of the IMU system (300Hz, see above) to obtain the number of spikes for each angular velocity value. We computed the head rotation modulatory index in freely moving mice as follow, hrMI = (movement FR - stationary FR) / (movement FR + stationary FR). Movement FR is the average firing rate between 70-90°/s (a velocity comparable to what has been used in head-fixed condition, i.e. 80°/s, see above). Stationary FR is the firing rate at 0°/s). To test the significance of the hrMI, we shuffled the spikes, computed the hrMI for 1000 randomizations and calculated the mean value *m* and standard deviation *σ* of obtained distribution of hrMIs. The statistical P-value of the hrMI measured in the experimental data set was computed as 1-F(hrMI, m, σ), where F is the cumulative Gaussian distribution with average *m* and standard deviation *σ*. Both fast spiking and regular spiking cells were merged together.

#### Cortical depth estimation

Cortical depth estimation from pia was performed using electrophysiological landmarks across layers. Average current source density (CSD) map and LFP traces during light flashes allows to locate layer 4 in V1^31^. Moreover, the Multi-unit (MUA) spectral power (500 Hz to 5 kHz) distribution along probe track allows to locate layer 5a^32^. Together, these two properties allowed us to normalize the cortical depth from the pia across mice.

### Statistics

Statistical analyses were done using Matlab. No statistical tests were used to predetermined sample size, but our sample sizes are similar to those generally employed in the field. All data are presented as mean ± standard error of the mean (SEM), unless otherwise noted. The stated P-values are the results of Wilcoxon rank sum test to compare values between different mice or recordings, and the Wilcoxon signed rank test to compare values from the same recording. Experiments and analyses were not blinded.

## Supplementary Figure Legends

**Supplementary Figure 1.**
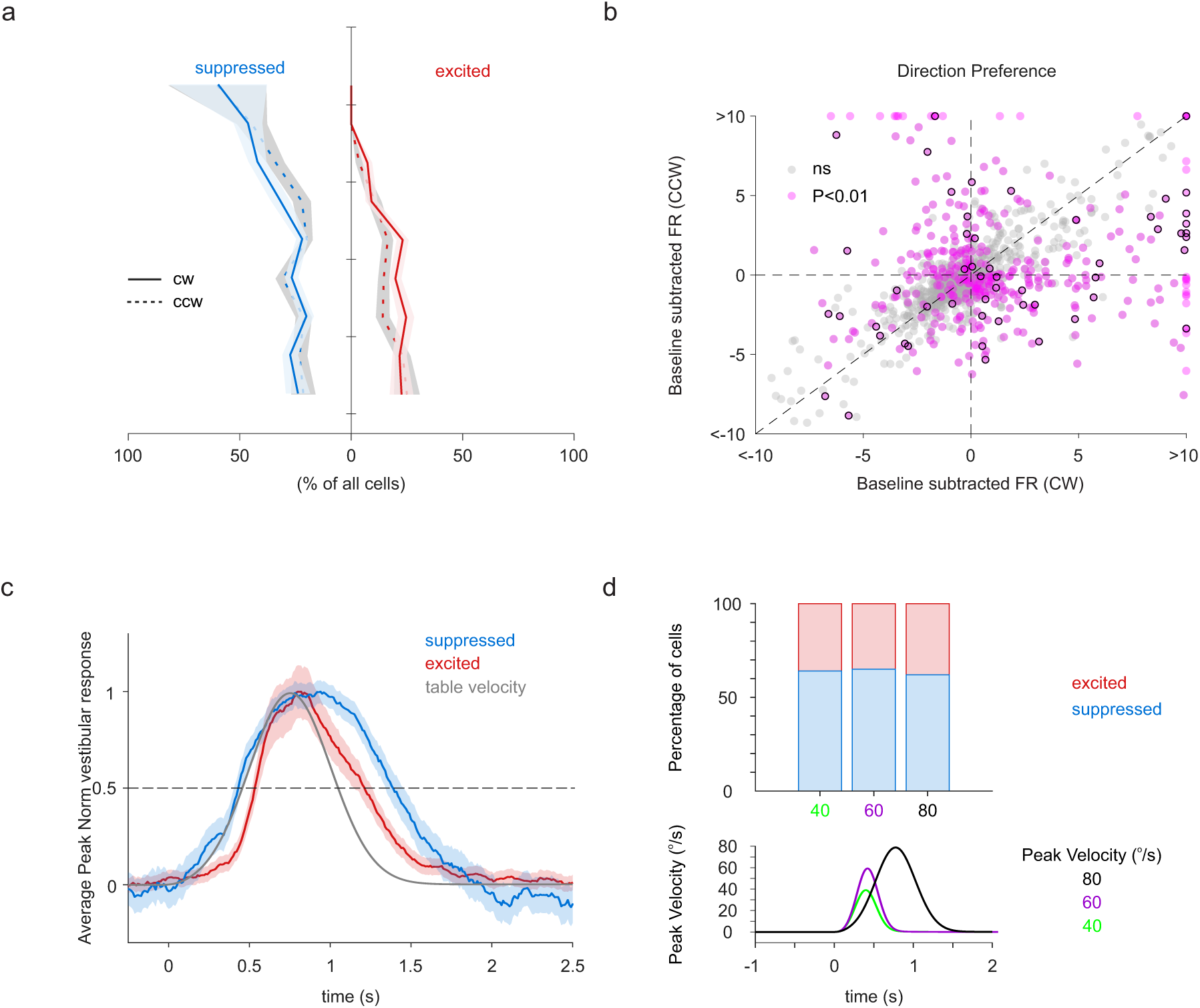
Impact of clockwise and counterclockwise head movements on V1 activity. **(a)** Percentage of suppressed (blue) and excited (red) cells plotted against cortical depth in response to CW (solid line) and CCW rotations (dashed line). Shading is SEM. Percentages of modulated units CW vs. CCW: 28±2.3 vs 28±1.8% suppressed; P = 0.90; 20±1.6 vs 17±1% excited; P = 0.18. Cortical depth is calculated from the pia. Median vMI: 0.388±0.01 and 0.387±0.01 for CW and CCW rotations, respectively (P = 0.25, n = 928 cells). **(b)** Baseline subtracted firing rate (FR) of individual cells in response to CW rotations plotted against the response to CCW rotations. Magenta data-points represent cells showing a significantly different (P<0.01) response between CW and CCW rotations. Magenta data-points with or without black border are FS and RS cells, respectively. Gray data points are cells showing no significant difference between their response to CW and CCW rotations. 62% of all cells (928 out of 1502 cells, n = 30 animals) was significantly modulated by either CW rotations (703 out of 1502 cells, 47%), CCW rotations (657 out of 1502 cells, 44%) or both (432 out of 1502 cells, 29%; CW only: 271 out of 1502 cells, 18%; CCW only: 225 out of 1502 cells, 15%). The magenta data points represent 40% of all significantly modulated cells. **(c)** Peak-normalized averaged modulation of suppressed (blue; n = 356 cells) and excited (red; n = 299 cells) cells across all layers in response to CW rotations. Gray line: velocity profile. Dotted line is half maximal amplitude. Half width at half maximal amplitude is 1 s and 0.69 s for suppression and excitation, respectively. **(d)** The overall suppressive impact of CW rotation on V1 activity does not depend on the specific velocity profile. Top panel: Percentage of significantly suppressed (blue) and excited cells for the three velocity profiles illustrated below (40°/s in green, 64% suppressed vs. 36% facilitated, n = 335 cells; 60°/s in purple, 65% suppressed vs. 35% facilitated, n = 335 cells; 80°/s in dark, 62% suppressed vs. 38% facilitated, n = 1502 cells).

**Supplementary Figure 2.**
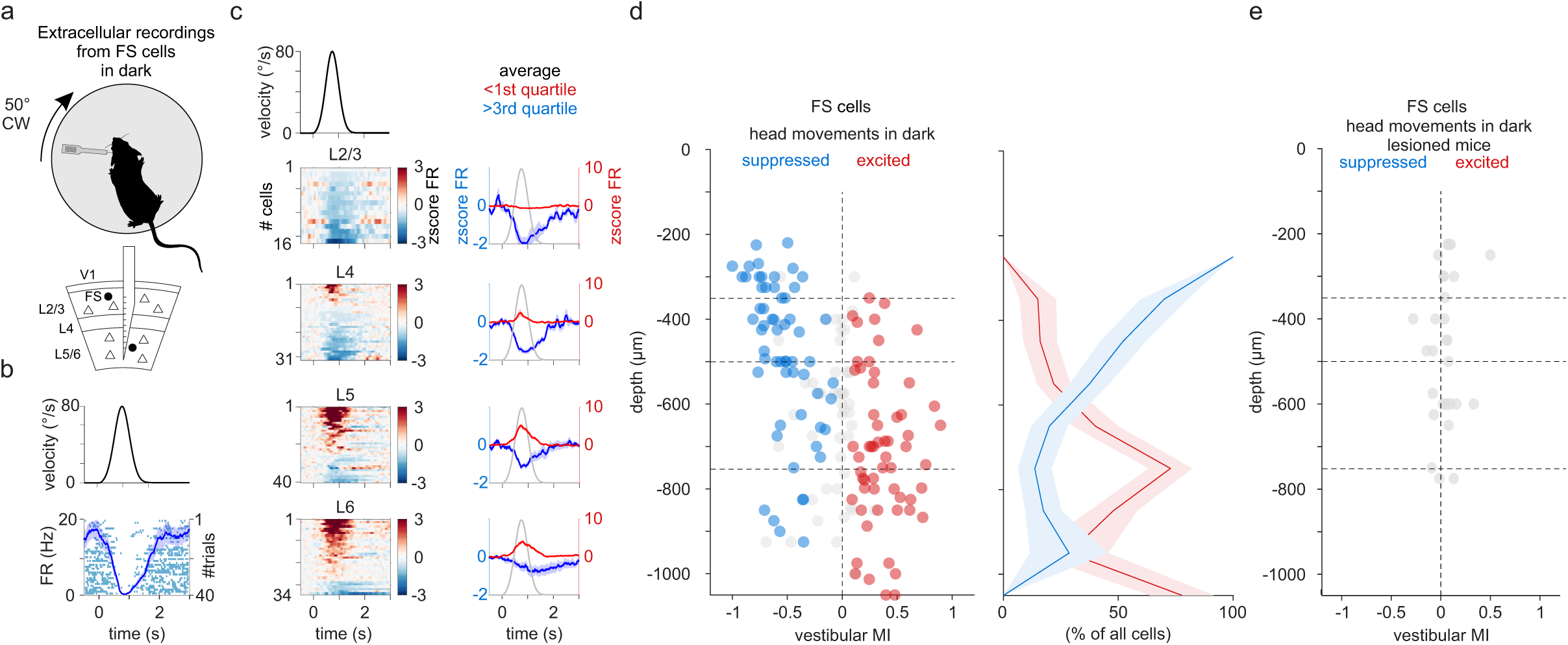
Suppression of FS cells by head movements in dark. **(a)** Experimental configuration: Top: An extracellular linear probe in the left primary visual cortex (V1) of a head-fixed, awake mouse records the response to clockwise (CW) rotations of the table in the dark. Bottom: The linear probe spanned all cortical layers. FS: fast spiking cell. **(b)** Top: Velocity profile of the rotating table (CW rotation). Bottom: Example FS cells located 725 μm below the pial surface. Raster plot and averaged peristimulus time histogram (PSTH) are superimposed. Shading is SEM. **(c)** Summary of 121 FS cells significantly modulated cells subdivided according to cortical depth (from the pia). Left panels: Velocity profile of the rotating table (CW rotation; top panel) and heat maps of the z-score of the firing rate (zscore FR) of individual FS cells during rotations. Within each panel, cells are sorted by the peak z-score. Right panels: Blue and red traces are the average z-scores of all cells whose z-score was smaller or larger than the first or third quartile, respectively. Shaded area is SEM. The gray trace is the velocity profile (CW rotation). **(d)** Left: vMI for CW rotations of individual FS cells plotted against cortical depth. Blue, red and gray circles are suppressed, excited and non-significantly modulated FS cells, respectively (Superficial layers vMI: −0.41±0.06; n = 51 cells; Deep layers vMI: 0.04±0.03; n = 112 cells; 36 recordings from 30 mice). Horizontal dotted lines indicate approximate layer borders. Right: Percentage of suppressed (blue) and excited (red) FS units plotted against cortical depth. Shading is SEM. **(e)** As (d) but for vestibular lesioned mice. (0 out of 29 FS cells were modulated, n = 3 recordings from 3 mice)

**Supplementary Figure 3.**
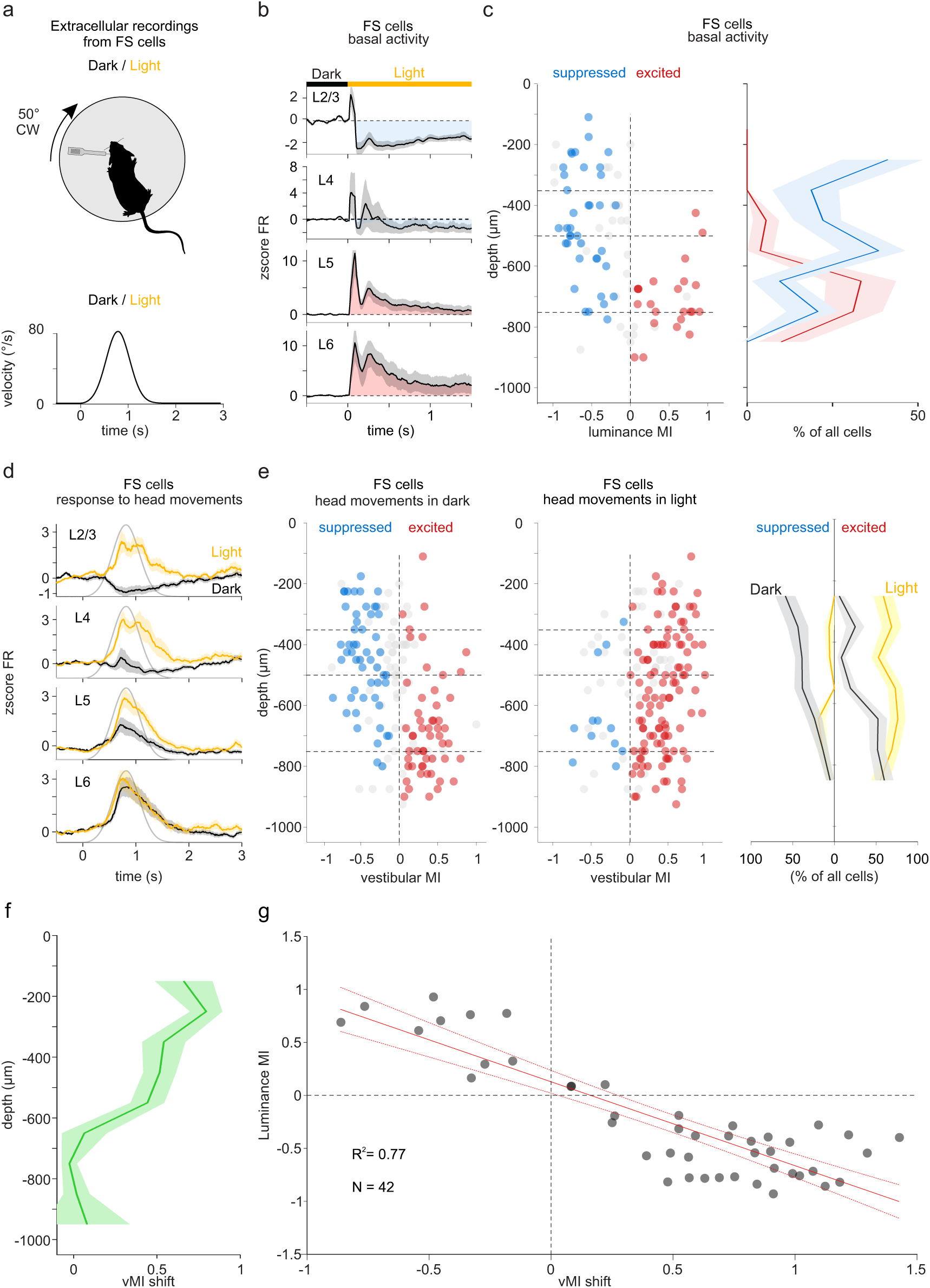
Excitation of FS cells by head movements in light. **(a)** Experimental configuration: Top: As in Supplementary Figure 2a, but this time rotations are alternated between dark and light. Bottom: Velocity profile of the rotation. **(b)** Averaged z-score of the firing rate (zscore FR) of FS cells in response to light onset. Each average is contributed by all the units recorded within the layer indicated in the panel, independently of whether they were excited, suppressed on non-significantly modulated by light onset. Gray shading is SEM. Blue or red shaded area indicate average suppression or facilitation by light, respectively. Horizontal bar indicates time of dark/light transition (time 0). **(c)** Luminance modulation index (lumMI) of individual FS cells plotted against cortical depth. Blue, red and gray circles represent FS cells whose basal activity is suppressed, excited and non-significantly modulated by light onset, respectively. Superficial FS cells lumMI: −0.44±0.05; Deep FS cells lumMI: −0.025±0.05. Horizontal dotted lines indicate approximate layer borders. Right: percentage of suppressed (blue) and excited (red) RS cells plotted against cortical depth in response to light onset. Shading is SEM. **(d)** Averaged z-score of the firing rate (zscore FR) of FS cells during CW rotations in dark (black traces) and in light (yellow traces). Each average is contributed by all the FS cells recorded within the layer indicated in the panel, independently of whether they were excited, suppressed on non-significantly modulated by the rotation. Shading is SEM. Gray trace is velocity profile. **(e)** Vestibular modulation index (vMI) for CW rotations of individual FS cells plotted against cortical depth. Blue, red and gray circles are suppressed, excited and non-significantly modulated FS cells, respectively. Same cells as in C. Left: vMI of FS cells in dark; Middle, vMI of the same FS cells in light (n = 170 RS cell; 18 recordings from 12 mice). Dark vs Light: Superficial layers vMI (n = 71 cells): −0.24±0.04 vs 0.35±0.04; P = 3.3e-10; Deep layers vMI (n = 99 cells): 0.098±0.037 vs 0.22±0.04; P = 0.04. Horizontal dotted lines indicate approximate layer borders. Right: percentage of suppressed and excited FS cells plotted against cortical depth in dark (black) and light (yellow). Shading is SEM. Note the shift towards positive vMI in light. **(f)** vMI shift (vMI in light - vMI in dark) plotted against cortical depth. Shading is SEM. Positive vMI shift indicate facilitation of the response to head movements in light as compared to dark. **(g)** vMI shift plotted against lumMI. Included are only FS cells with a significant lumMI and a significant vMI shift (n = 42). Continuous red line: linear regression (R^2^ = 0.77). Dotted red lines: confidence interval of linear regression. Note inverse relationship between vMI shift and lumMI.

**Supplementary Figure 4.**
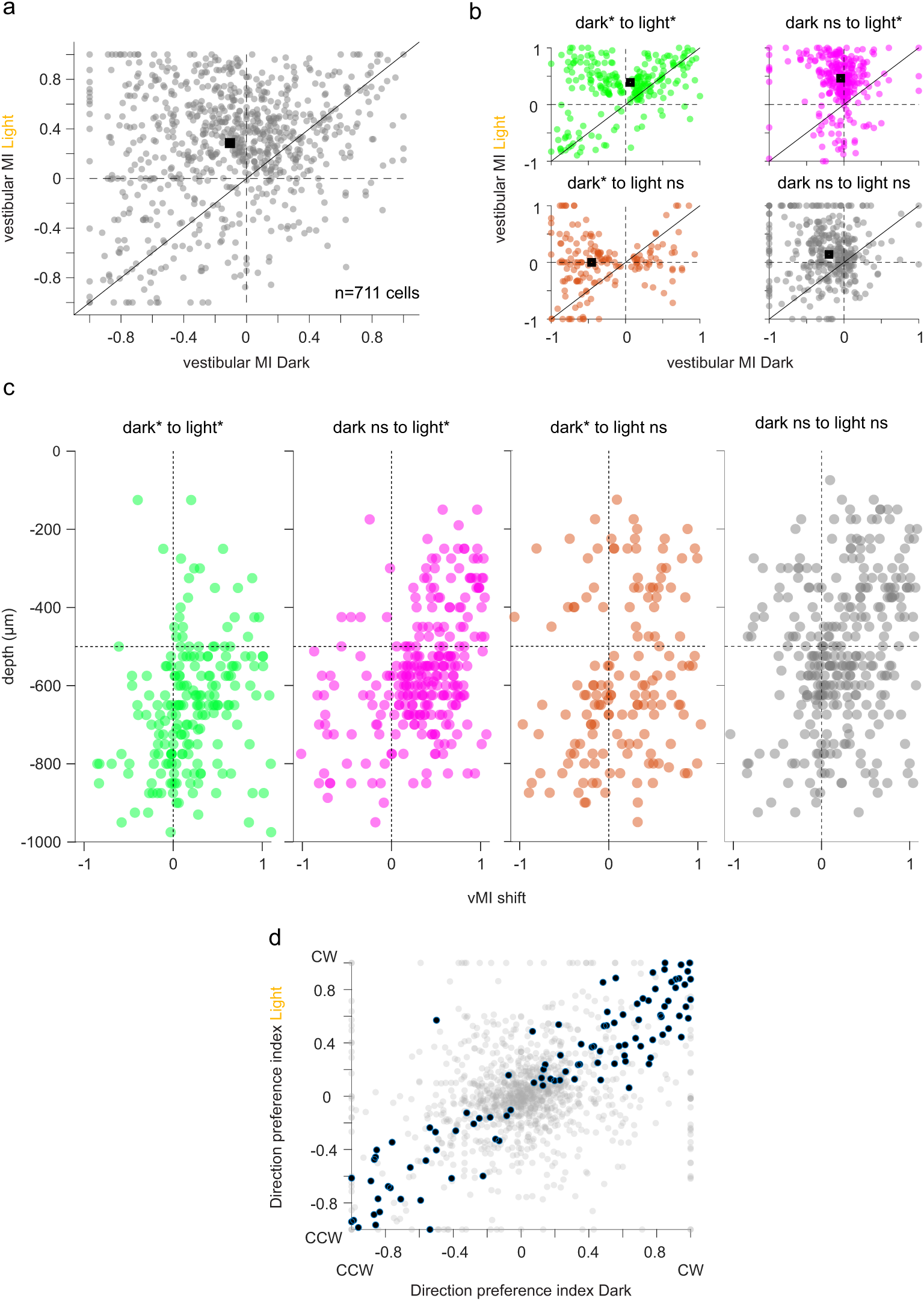
Comparing responses to head movements in dark versus light for individual RS cells. **(a)** The vestibular modulation index (vMI) to CW rotations in dark is plotted against the vMI in light for individual RS cells that are significantly modulated by head movements in either dark or light (n = 711 cells). The dark square is the median. **(b)** Same as in (a) but after selecting RS cells according to four different criteria: Green: vMI is significant (P<0.05) both in dark an in light (n = 235 cells). Magenta: vMI is significant in light but not in dark (n = 296 cells). Orange: vMI is significant in dark but not in light (n = 180 cells). Gray: vMI is not significant in either condition (n = 376 cells; note that cells in gray are not included in the plot in (A)). The dark square is the median. dark*: vMI is significant in dark. light*: vMI is significant in light; dark ns: vMI is non-significant in dark; light ns: vMI is non-significant in light. **(c)** vMI shift (vMI in light - vMI in dark) plotted against cortical depth. Same cells and color-code as in (B). Cortical depth is measured from the pia. **(d)** Direction preference index (see methods) in dark plotted against direction preference in light for RS cells (n = 1087 cells; same cells shown in (b)). Dark data-points represent cells showing a significantly different (P<0.01) response between CW and CCW rotations in both dark and light conditions (n = 103 cells). Note that negative values represent a preference for CCW rotation whereas positive values represent a preference for CW rotations.

**Supplementary Figure 5.**
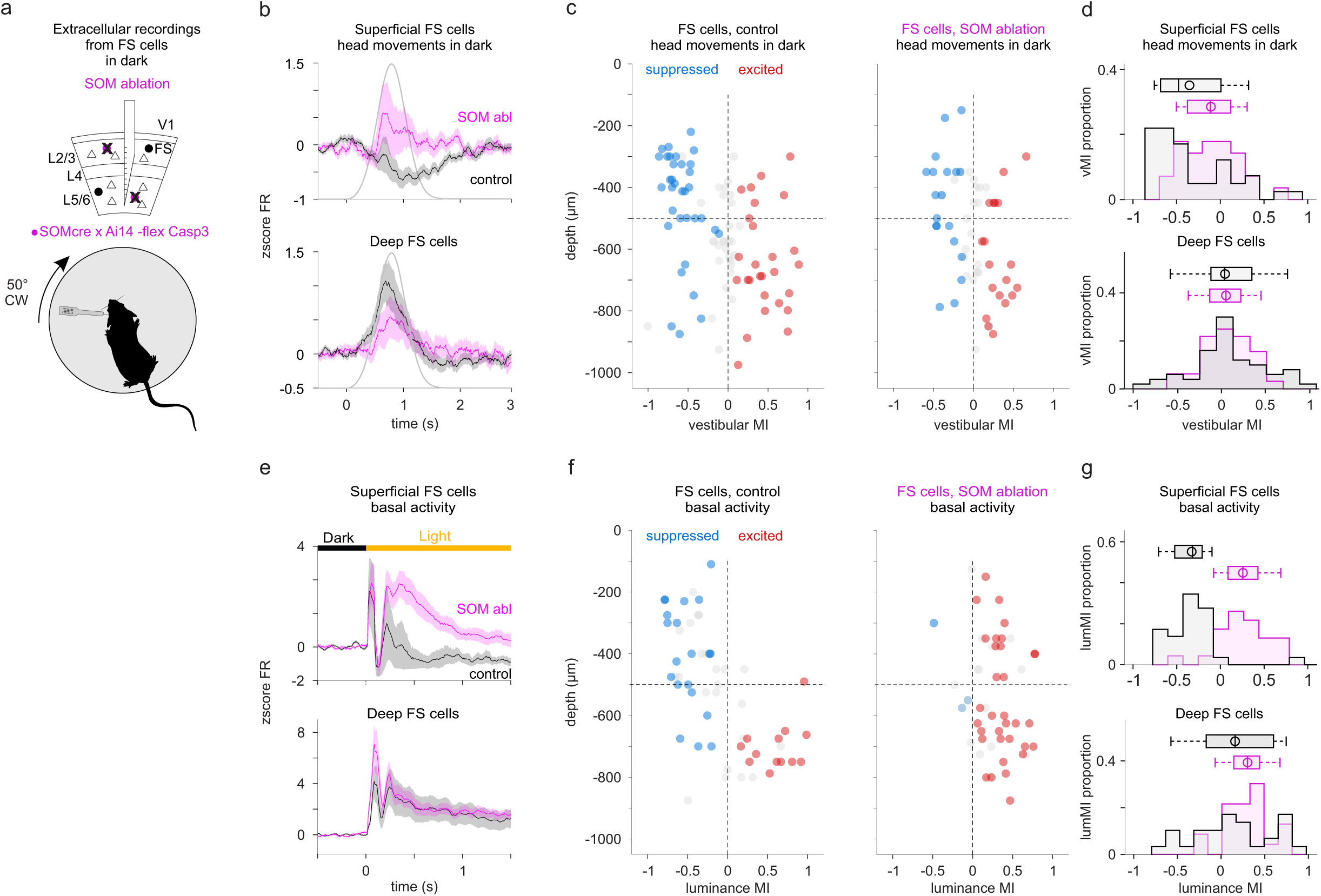
SOM cells suppress FS cells in response to head movements and light onset. **(a)** Experimental configuration: Top: Schematic of extracellular recordings from V1 in which SOM cells have been ablated. Bottom: The extracellular linear probe is inserted in the left V1 of the head-fixed, awake mouse to record the activity of FS units in response to clockwise (CW) rotation of the table in dark. **(b)** Averaged z-score of the firing rate (zscore FR) of FS cells during CW rotations in dark for SOM ablated animals (magenta) and un-injected, control littermates (black). Top panel: All FS cells recorded in superficial layers (n = 26 cells (magenta) and 37 cells (black)). Bottom panel: All FS cells recorded in deep layers (n = 34 cells (magenta) and 54 cells (black)). Shading is SEM. Gray trace is velocity profile. **(c)** Vestibular modulation index (vMI) for CW rotations of individual FS cells plotted against cortical depth. Blue, red and gray circles are suppressed, excited and non-significantly modulated FS cells, respectively. Left: vMI of FS cells in un-injected littermates (n = 91 FS cell; 11 mice; 1 recording per animal). Right: vMI of FS units in SOM ablated animals (n = 60 FS cell; 9 mice; 1 recording per animal). Control vs. ablation: Superficial FS cells vMI: −0.33±0.07 (n = 37) vs −0.045±0.06 (n = 26); P = 0.003; Deep FS cells vMI: 0.044±0.06 (n = 54) vs 0.03±0.05 (n = 34); P = 0.94. Horizontal dotted line indicates approximate border between superficial and deep layers (between layer 4 and 5). Note shift towards positive vMI of FS cells in superficial layers of SOM ablated animals. **(d)** Distribution of the vMI in SOM ablated animals (magenta) and control littermates (black) for FS cells recorded in superficial (top panel) and deep (bottom panel) layers. Box plots show median, first and third quartiles, minimum/maximum values and mean (circle). **(e)** Averaged z-score of the firing rate (zscore FR) of FS cells in response to light onset in SOM ablated animals (magenta) and control littermates (black). Top panel: FS cells recorded in superficial layers (n = 26 cells (magenta) and 37 cells (black)). Bottom panel: FS cells recorded in deep layers (n = 34 cells (magenta) and 54 cells (black)). Shading is SEM. Horizontal bar indicates time of dark/light transition (time 0). **(f)** Luminance modulation index (lumMI) of individual FS cells plotted against cortical depth. Blue, red and gray circles are suppressed, excited and non-significantly modulated FS cells, respectively. Left: lumMI of FS cells in un-injected littermates (n = 58 FS cell; 8 mice; 1 recording per animal); Right, vMI of SOM ablated animals (n = 46 FS cell; 8 mice; 1 recording per animal). Control vs Ablation: Superficial FS cells vMI: −0.32±0.06 (n = 27 cells) vs 0.25±0.06 (n = 21 cells); P = 5.5e-07; Deep FS cells vMI: 0.09±0.09 (n = 31 cells) vs 0.29±0.05 (n = 25 cells); P = 0.12. Note shift towards positive lumMI in SOM ablated animals. Horizontal dotted line indicates approximate border between superficial and deep layers. **(g)** Distribution of the lumMI in SOM ablated animals (magenta) and control littermates (black) for FS cells recorded in superficial (top panel) and deep (bottom panel) layers. Box plots show median, first and third quartiles, minimum/maximum values and mean (circle).

**Supplementary Figure 6.**
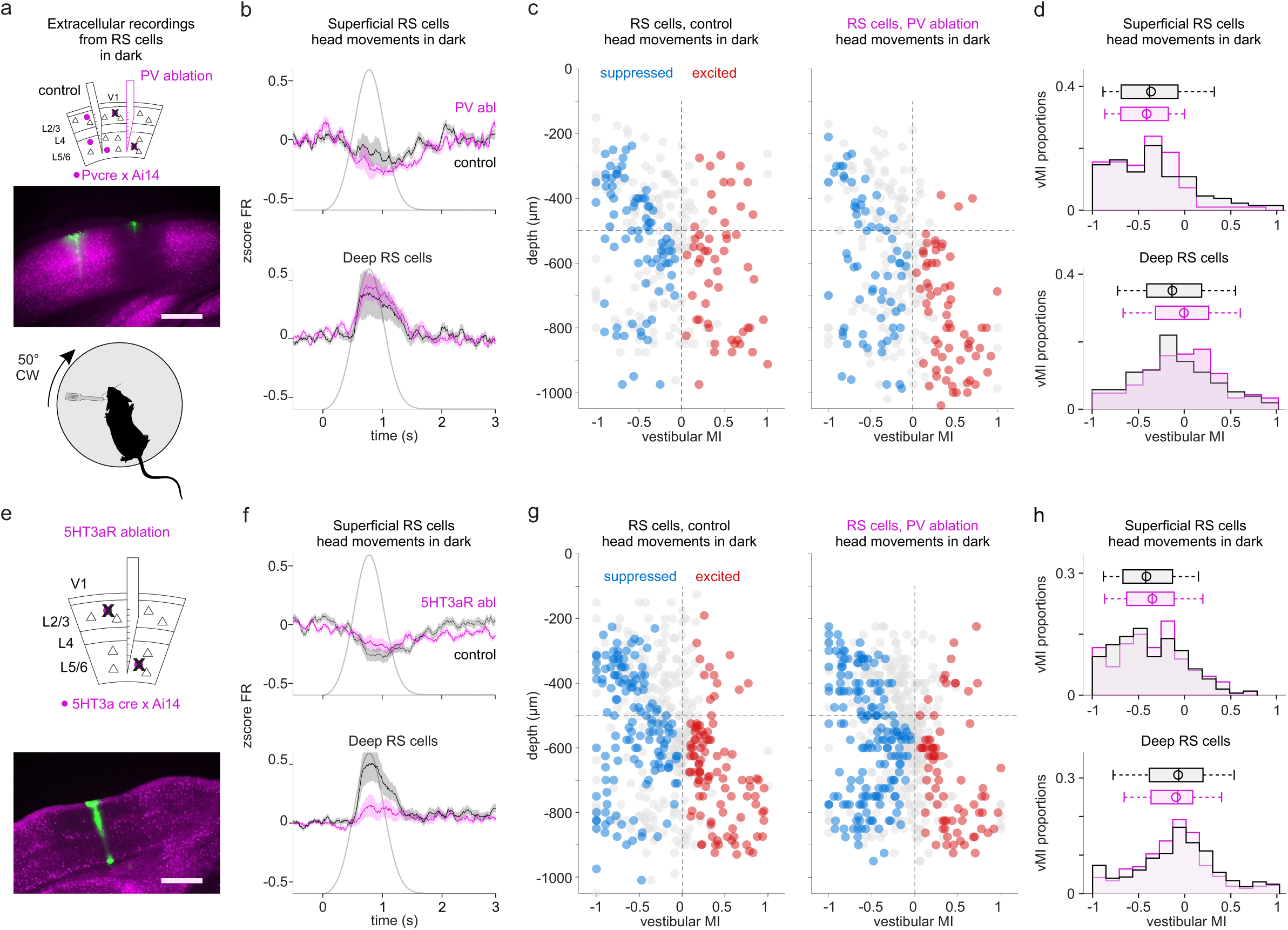
PV and 5HT3aR cells do not contribute to the suppression of V1 by head movements. **(a)** Experimental configuration: Top: Schematic of extracellular recordings from a site in V1 where PV cells have been ablated and from a neighboring, control site where PV cells are intact. Middle: Fluorescence microscopy image of a V1 coronal section (PV Cre x Ai14 mouse) with the conditional expression of virally injected caspase-3. The electrode tracks are in green. Note the reduction in fluorescence around the injection site as compared to the neighboring non-injected region. Scale bar: 400 μm. Bottom: Extracellular linear probe inserted in the left V1 of the head-fixed, awake mouse to record the activity of RS cells in response to clockwise (CW) rotation of the table in dark. **(b)** Averaged z-score of the firing rate (zscore FR) of RS cells during CW rotations in dark for PV ablated (magenta) and control sites (black). Top panel: All RS units recorded in superficial layers (n = 87 cells (magenta) and 122 cells (black)). Bottom panel: All RS cells recorded in deep layers (n = 206 cells (magenta) and 163 cells (black)). Shading is SEM. Gray trace is velocity profile of the rotating table (CW rotation). **(c)** Vestibular modulation index (vMI) for CW rotations of individual RS cells plotted against cortical depth. Blue, red and gray circles are suppressed, excited and non-significantly modulated RS cells, respectively. Left: vMI of RS cells in non-injected site (n = 285 RS cells; 5 mice). Right: vMI of RS cells in PV ablated site (n = 293 RS cells; 5 mice). Control vs PV cells ablation: Superficial RS cells vMI: −0.38±0.04 (n = 122 cells) vs −0.43±0.04 (n = 87 cells); P = 0.36; Deep RS cells vMI: −0.11±0.03 (n = 163 cells) vs −0.04±0.03 (n = 206 cells); P = 0.09. Horizontal dotted line indicates approximate border between superficial and deep layers (between layer 4 and 5). **(d)** Distribution of the vMI in PV ablated (magenta) and control sites (black) for RS cells recorded in superficial (top panel) and deep (bottom panel) layers. Box plots show median, first and third quartiles, minimum/maximum values and mean (circle). **(e)** Experimental configuration: Top: Schematic of extracellular recordings from a V1 where 5HT3aR cells have been ablated. Middle: Fluorescence microscopy image of a V1 coronal section (5HT3aR Cre x Ai14 mouse) with the conditional expression of virally injected caspase-3. The electrode track is in green. Note the reduction in fluorescence around the injection site. Scale bar: 400 μm. **(f)** Averaged z-score of the firing rate (zscore FR) of RS cells during CW rotations in dark for 5HT3aR ablated animals (magenta) and non-injected, control littermates (black). Top panel: All RS cells recorded in superficial layers (n = 190 cells (magenta) and 186 cells (black)). Bottom panel: All RS cells recorded in deep layers (n = 383 cells (magenta) and 335 cells (black)). Shading is SEM. Gray trace is velocity profile of the rotating table (CW rotation). **(g)** Vestibular modulation index (vMI) for CW rotations of individual RS cells plotted against cortical depth. Blue, red and gray circles are suppressed, excited and non-significantly modulated RS cells, respectively. Left: vMI of RS cells in non-injected littermates (n = 421 RS cells; 11 mice; 13 recordings). Right: vMI of RS cells in 5HT3aR ablated animals (n = 573 RS cell; 8 mice; 11 recordings). Control vs Ablation: Superficial RS cells vMI: −0.40±0.03 (n = 186 cells) vs −0.36±0.03 (n = 190); P = 0.36; Deep RS cells vMI: −0.09±0.02 (n = 335 cells) vs −0.12±0.02 (n = 383 cells); P = 0.17. Horizontal dotted line indicates approximate border between superficial and deep layers (between layer 4 and 5). **(h)** Distribution of the vMI in 5HT3aR ablated animals (magenta) and control littermates (black) for RS cells recorded in superficial (top panel) and deep (bottom panel) layers. Box plots show median, first and third quartiles, minimum/maximum values and mean (circle).

## Bibliography

1. Angelaki, D. E. Eyes on target: What neurons must do for the vestibulo-ocular reflex during linear motion. J. Neurophysiol. 92, 20–35 (2004).

2. Angelaki, D. E. & Cullen, K. E. Vestibular System: The Many Facets of a Multimodal Sense. Annu. Rev. Neurosci. 31, 125–150 (2008).

3. Rancz, E. et al. Widespread Vestibular Activation of the Rodent Cortex. J. Neurosci. 35, 5926–5934 (2015).

4. Vélez-Fort, M. et al. A Circuit for Integration of Head- and Visual-Motion Signals in Layer 6 of Mouse Primary Visual Cortex. Neuron 98, 179–191.e6 (2018).

5. Magnin, G. V.-M. and M. Single Neuron Activity Related to Natural Vestibular Stimulation in the Cat’s Visual Cortex. Exp. Brain Res. 451–455 (1982).

6. Duffy, C. J. MST neurons respond to optic flow and translational movement. J. Neurophysiol. 80, 1816–1827 (1998).

7. Laurens, J. et al. Transformation of spatiotemporal dynamics in the macaque vestibular system from otolith afferents to cortex. Elife 6, 1–26 (2017).

8. Bense, S., Stephan, T., Yousry, T. A., Brandt, T. & Dieterich, M. Multisensory cortical signal increases and decreases during vestibular galvanic stimulation (fMRI). J. Neurophysiol. 85, 886–899 (2001).

9. Wenzel, R. et al. Deactivation of human visual cortex during involuntary ocular oscillations A PET activation study. Brain 119, 101–110 (1996).

10. Gu, Y., Watkins, P. V., Angelaki, D. E. & DeAngelis, G. C. Visual and nonvisual contributions to three-dimensional heading selectivity in the medial superior temporal area. J. Neurosci. 26, 73–85 (2006).

11. Ohshiro, T., Angelaki, D. E. & DeAngelis, G. C. A Neural Signature of Divisive Normalization at the Level of Multisensory Integration in Primate Cortex. Neuron 95, 399–411.e8 (2017).

12. Deneux, T. et al. Context-dependent signaling of coincident auditory and visual events in primary visual cortex. Elife 8, 1–23 (2019).

13. Iurilli, G. et al. Sound-Driven Synaptic Inhibition in Primary Visual Cortex. Neuron 73, 814– 828 (2012).

14. Kinoshita, M. & Komatsu, H. Neural representation of the luminance and brightness of a uniform surface in the macaque primary visual cortex. J. Neurophysiol. 86, 2559–2570 (2001).

15. Kayama, Y., Riso, R. R., Bartlett, J. R. & Doty, R. W. Luxotonic responses of units in macaque striate cortex. J. Neurophysiol. 42, 1495–1517 (1979).

16. Tucker, T. R. & Fitzpatrick, D. Luminance-Evoked Inhibition in Primary Visual Cortex: A Transient Veto of Simultaneous and Ongoing Response. J. Neurosci. 26, 13537–13547 (2006).

17. Xing, D., Yeh, C.-I., Gordon, J. & Shapley, R. M. Cortical brightness adaptation when darkness and brightness produce different dynamical states in the visual cortex. Proc. Natl. Acad. Sci. 111, 1210–1215 (2014).

18. Pfeffer, C. K., Xue, M., He, M., Huang, Z. J. & Scanziani, M. Inhibition of inhibition in visual cortex: the logic of connections between molecularly distinct interneurons. Nat. Neurosci. 16, 1068–76 (2013).

19. Bortone, D. S., Olsen, S. R. & Scanziani, M. Translaminar inhibitory cells recruited by layer 6 corticothalamic neurons suppress visual cortex. Neuron 82, 474–485 (2014).

20. Lima, S. Q., Hromádka, T., Znamenskiy, P. & Zador, A. M. PINP: A new method of tagging neuronal populations for identification during in vivo electrophysiological recording. PLoS One 4, (2009).

21. Lee, S. H., Hjerling-Leffler, J., Zagha, E., Fishell, G. & Rudy, B. The largest group of superficial neocortical GABAergic interneurons expresses ionotropic serotonin receptors. J. Neurosci. 30, 16796–16808 (2010).

22. Ma, Y., Hu, H., Berrebi, A. S., Mathers, P. H. & Agmon, A. Distinct subtypes of somatostatin-containing neocortical interneurons revealed in transgenic mice. J. Neurosci. 26, 5069–5082 (2006).

23. Nigro, M. J., Hashikawa-Yamasaki, Y. & Rudy, B. Diversity and connectivity of layer 5 somatostatin-expressing interneurons in the mouse barrel cortex. J. Neurosci. 38, 1622– 1633 (2018).

24. Naka, A. et al. Complementary networks of cortical somatostatin interneurons enforce layer specific control. Elife doi:10.7554/eLife.43696.

25. Wang, Q. & Burkhalter, A. Area map of mouse visual cortex. J. Comp. Neurol. 502, 339– 357 (2007).

26. Leinweber, M., Ward, D. R., Sobczak, J. M., Attinger, A. & Keller, G. B. A Sensorimotor Circuit in Mouse Cortex for Visual Flow Predictions. Neuron 95, 1420–1432.e5 (2017).

27. Storchi, R. et al. Modulation of Fast Narrowband Oscillations in the Mouse Retina and dLGN According to Background Light Intensity. Neuron 93, 299–307 (2017).

28. Ji, X. Y. et al. Thalamocortical Innervation Pattern in Mouse Auditory and Visual Cortex: Laminar and Cell-Type Specificity. Cereb. Cortex 26, 2612–2625 (2016).

29. Silberberg, G. & Markram, H. Disynaptic Inhibition between Neocortical Pyramidal Cells Mediated by Martinotti Cells. Neuron 53, 735–746 (2007).

30. Pasquet, M. O. et al. Wireless inertial measurement of head kinematics in freely-moving rats. Sci. Rep. 6, 1–13 (2016).

31. Niell, C. M. & Stryker, M. P. Highly selective receptive fields in mouse visual cortex. J. Neurosci. 28, 7520–7536 (2008).

32. Senzai, Y., Fernandez-Ruiz, A. & Buzsáki, G. Layer-Specific Physiological Features and Interlaminar Interactions in the Primary Visual Cortex of the Mouse. Neuron 101, 500–513.e5 (2019).

